# Lewis rat NLRP1 inflammasome activation is mediated by three *Toxoplasma gondii* dense granule proteins

**DOI:** 10.1101/381202

**Authors:** Yifan Wang, Kimberly M. Cirelli, Patricio D.C. Barros, Lamba Omar Sangaré, Vincent Butty, Musa A. Hassan, Patricia Pesavento, Asli Mete, Jeroen P.J. Saeij

**Author notes:** Present address: Kimberly M. Cirelli, Division of Vaccine Discovery, La Jolla Institute for Allergy and Immunology, La Jolla, California 92037, USA. Present address: Patricio D.C. Barros, Laboratory of Immunoparasitology “Dr. Mário Endsfeldz Camargo”, Department of Immunology, Institute of Biomedical Sciences, Federal University of Uberlândia, Uberlândia, Brazil. Y.W. and K.M.C. contributed equally to this work. Address correspondence to Jeroen P.J. Saeij.

## Abstract

The Lewis rat is the only known warm-blooded animal that has sterile immunity to *Toxoplasma*. Upon invasion of Lewis rat macrophages *Toxoplasma* rapidly activates the nucleotide-binding oligomerization domain, leucine-rich repeat and pyrin domain containing 1 (NLRP1) inflammasome resulting in interleukin (IL)-1β secretion and a form of cell death known as pyroptosis, which prevents *Toxoplasma* replication. Using a chemical mutagenesis screen we identified *Toxoplasma* mutants that no longer induced pyroptosis. Whole genome sequencing led to the identification of three *Toxoplasma* parasitophorous vacuole-localized dense granule proteins, GRA35, GRA42 and GRA43 that are individually required for inflammasome activation in Lewis rat macrophages. Macrophage infection with Δ*gra35*, Δ*gra42*, and Δ*gra43* parasites leads to greatly reduced cell death and reduced IL-1β secretion. Lewis rat macrophage infected with parasites containing single, double or triple deletion of these GRAs showed similar levels of cell viability suggesting the three GRAs function in the same pathway that activates the inflammasome. Deletion of *GRA42* and *GRA43* resulted in GRA35, and other GRAs, being retained inside the parasitophorous vacuole instead of being localized to the parasitophorous vacuole membrane. *Toxoplasma* deficient in GRA35, GRA42 or GRA43 do not establish chronic infection in Lewis rats, but have reduced cyst number in parasite-susceptible F344 rats, in which *Toxoplasma* does not activate the NLRP1 inflammasome, revealing these GRAs determine parasite *in vivo* fitness independent of their role in inflammasome activation. Overall, our data suggest that *Toxoplasma* dense granule proteins that localize to the parasitophorous vacuole membrane are novel mediators of host NLRP1 inflammasome activation.

**Importance:** Inflammasomes are a major component of the innate immune system and responsible for detecting various microbial and environmental danger signals. The Lewis rat has sterile immunity to *Toxoplasma* because upon invasion of Lewis rat macrophages the parasite rapidly activates the NLRP1 inflammasome resulting in cell death and parasite elimination. The work reported here identified that *Toxoplasma* GRA35, GRA42 and GRA43 are required for activation of the Lewis rat NLRP1 inflammasome. GRA42 and GRA43 mediate the correct localization of other GRAs, including GRA35, to the parasitophorous vacuole membrane. In addition to their role in inflammasome activation, these three GRAs are also important for parasite *in vivo* fitness in a *Toxoplasma*-susceptible rat strain. Thus, these results give new insight into NLRP1 inflammasome activation by *Toxoplasma* effectors and identified three GRAs that are required for pathogenesis of the parasite.

## Introduction

*Toxoplasma* is an obligate intracellular protozoan parasite that infects a wide variety of warm-blooded animals (1). Among its different hosts there are natural differences in susceptibility to the parasite. Most laboratory mouse strains are susceptible to infection and can succumb after low dose injection of virulent parasite strains. Rats and humans are relatively resistant to *Toxoplasma*. Most rat strains remain asymptomatic after infection, but the parasite establishes a chronic infection by developing into cysts in brain and muscle tissues. However, the Lewis rat strain can clear the parasite and fails to develop a chronic infection (2). This resistance was shown to be a myeloid cell-intrinsic dominant trait that mapped to a single locus, *Toxo1* (3). *In vitro*, resistance correlates with rapid induction of Lewis rat macrophage cell death after *Toxoplasma* invasion (4–6). *Toxoplasma*-induced Lewis macrophage cell death is controlled by *Nlrp1*, which encodes for the NLRP1 inflammasome sensor (4, 5).

The inflammasomes are a family of cytosolic pattern recognition receptors (PRRs). Activation of the sensor, leads to the formation of a multimeric complex and the recruitment and proteolytic activation of pro-caspase-1. Caspase-1 cleaves the cytokines pro-IL-1β and pro-IL-18 resulting in their release from the cells. Caspase-1 activation also cleaves Gasdermin D (GSDMD) which can subsequently form pores in the host cell membrane and is therefore an essential trigger for a type of host cell death, termed pyroptosis (7, 8). Pyroptosis is a highly inflammatory form of programmed cell death that occurs most frequently upon infection with intracellular pathogens and has been established as a host mechanism to clear intracellular pathogens (9). *Toxoplasma* infection in Lewis rat bone marrow-derived macrophages (BMDMs) leads to NLRP1 inflammasome activation, which results in the release of IL-1β and IL-18 and pyroptosis of infected BMDMs, releasing parasites into the extracellular space before replication can occur (4, 5). As macrophages are among the predominant cell type infected upon *Toxoplasma* infection (10), it is likely that macrophage pyroptosis is a host mechanism to prevent parasite proliferation inside the host. Furthermore, infected macrophages and dendritic cells are involved in promoting *Toxoplasma* dissemination by migrating to distant sites (11–13), and therefore *Toxoplasma*-induced pyroptosis of these cells could also inhibit *Toxoplasma* dissemination.

The specific stimuli that can activate the inflammasomes and their mechanism of activation vary. NLR family CARD domain-containing protein 4 (NLRC4) recognizes NLR family, apoptosis inhibitory protein (NAIP) proteins bound to bacterial components, namely flagellin and type III secretory system proteins (14). The NLRP3 inflammasome is activated by a large number of stimuli, such as low intracellular potassium concentrations (15), viruses e.g. influenza A (16), bacterial toxins e.g. nigericin and maitotoxin (17) and parasites e.g. *Plasmodium-derived* hemozoin (18). Anthrax Lethal Toxin (LT) is a protease and a direct activator of rat NLRP1 (19). LT cleaves the N-terminus of NLRP1 in LT-susceptible mouse and rat macrophages. This cleavage is sufficient to activate the NLRP1 inflammasome and induce pyroptosis (20). Inflammasome activation by *Toxoplasma* in mice was also recently evaluated (6, 21). No cleavage of the mouse NLRP1 was observed in parasite-infected cells suggesting the NLRP1 response to *Toxoplasma* in mice might be cleavage-independent (6). However, the parasite effector(s) that activate the NLRP1 inflammasome are unknown.

To further explore the mechanism of activation of the Lewis rat NLRP1 inflammasome by *Toxoplasma*, we chose to take an unbiased approach to identify the *Toxoplasma* gene product(s) required for activation of Lewis rat BMDM cell death. Using a chemical mutagenesis screen followed by whole genome sequencing we identified three *Toxoplasma* dense granule proteins (GRA35, GRA42 and GRA43) that are required for inflammasome activation in Lewis rat macrophages. Parasite strains deficient in GRA35, GRA42 or GRA43 induce significantly less pyroptosis and IL-1β processing and secretion. These results indicate that *Toxoplasma* dense granule proteins are novel mediators of NLRP1 inflammasome activation.

## Results

### The NLRP3 inflammasome is not involved in *Toxoplasma*-induced Lewis rat macrophage cell death

We previously showed that *Toxoplasma* activates the NLRP1 inflammasome in Lewis rat macrophages resulting in rapid cell death (5). *Toxoplasma* activates both the NLRP1 and NLRP3 inflammasomes in mice (21) but it is not known whether *Toxoplasma* also activates the NLRP3 inflammasome in Lewis rat macrophages. To investigate this, Lewis rat macrophages were treated with NLRP3 inflammasome inhibitor MCC950 (22) or with the Caspase-1 inhibitor VX765 (which should inhibit all inflammasomes) (23) followed by *Toxoplasma* type I (RH) parasite infection. Infected macrophages treated with VX765 showed significantly higher cell viability compared to non-treated macrophages, whereas treatment with MCC950 did not prevent parasite-induced cell death (**Figure 1**). VX765 and MCC950 did not inhibit parasite invasion in Lewis rat macrophages (**Figure S1A**) nor parasite growth in HFFs (**Figure S1B**). As a positive control, MCC950 inhibited cell death and IL-1β release in response to Nigericin, a known NLRP3 agonist, in lipopolysaccharide (LPS)-primed Lewis rat macrophages (**Figure S1C and D**). Therefore, Lewis rat macrophage cell death upon *Toxoplasma* infection is likely entirely dependent on NLRP1.

**Figure 1.**
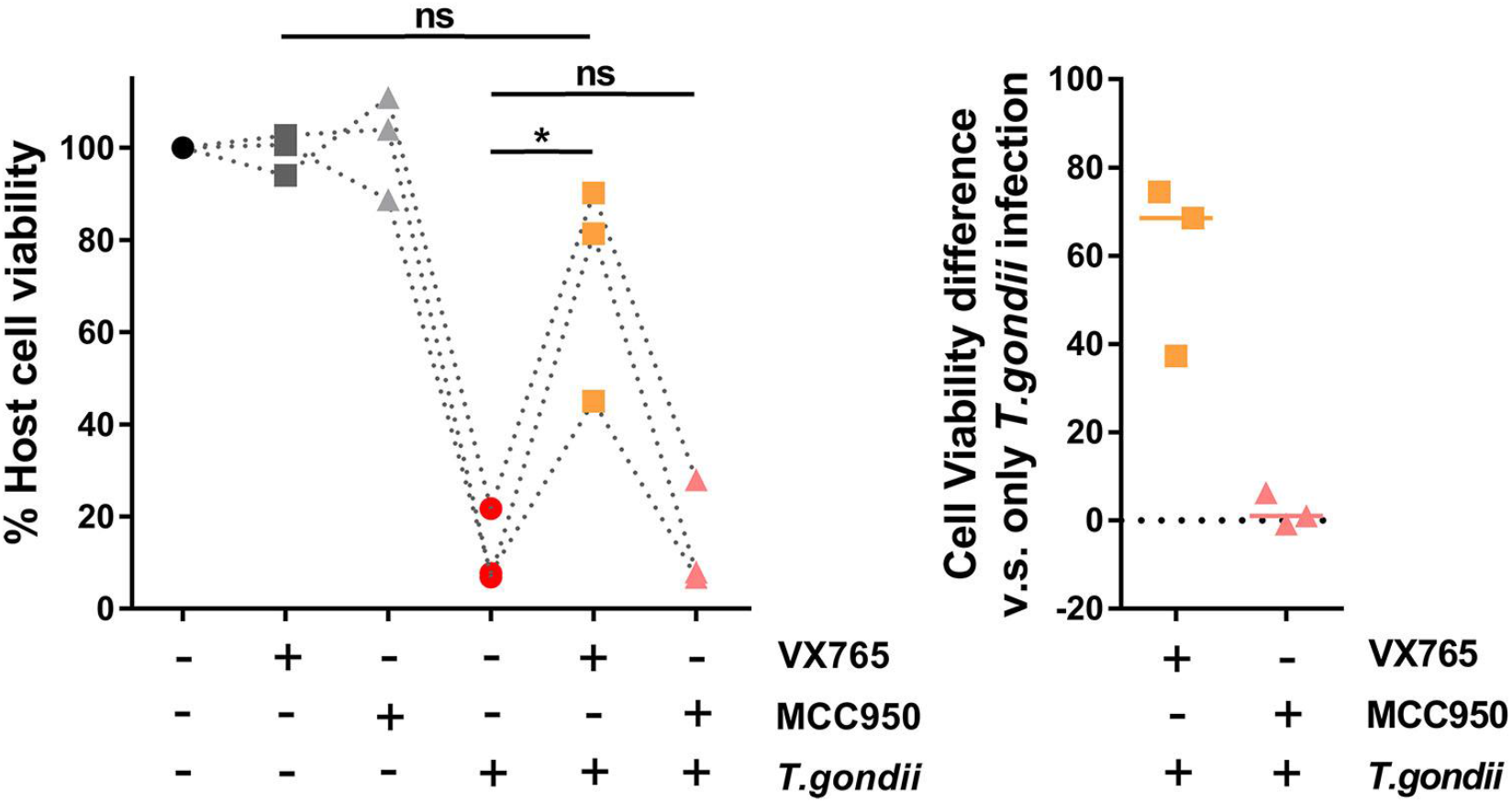
The NLRP3 inflammasome is dispensable for *Toxoplasma*-induced Lewis rat macrophage cell death and IL-1β secretion. Lewis rat BMDMs with or without pre-treatment of either 50 μM of VX765 or 10 μM of MCC950 for 2 hours were infection with *Toxoplasma* type I (RH) parasites (MOI = 0.5) for 24 hours. Macrophage viability was measured via 3-(4,5-dimethylthiazol-3-yl)-5-(3-carboxymethoxyphenyl)-2-(4-sulfophenyl)-2H-tetrazolium) (MTS) assay. Data are displayed as the paired scatterplots (left, *n* = 3; \**p* < 0.05, ns, not significant; student’s t-test). The right scatterplots are showing the cell viability difference between infected BMDMs with and without treatment in each paired experiment. Horizontal bars represent the median cell viability difference.

### *Toxoplasma*-induced Lewis rat macrophage cell death is dependent on Golgi-protease ASP5 but not MYR1

To better understand the mechanism of NLRP1 inflammasome activation, we aimed to discover the *Toxoplasma* effector(s) that mediate the activation of the Lewis rat NLRP1 inflammasome. We focused on parasite secretory proteins that can potentially interact with host cytosolic NLRP1 or interact with other host cytosolic proteins that modulate the activity of the inflammasome. Upon invasion, *Toxoplasma* secretes rhoptry proteins (ROPs) into the host cell cytosol (24). We previously showed that parasites treated with Mycalolide B, a compound that blocks *Toxoplasma* invasion but allows for secretion of microneme and rhoptry contents, were unable to induce Lewis rat macrophage IL-1β secretion and cell death (5) suggesting that ROPs are not the parasite effectors that activate the NLRP1 inflammasome. Once the parasite resides inside a host cell in a non-fusogenic parasitophorous vacuole (PV), dense granules discharge GRAs into the PV lumen, associated with the PV membrane (PVM), or are exported into the host cytosol (25). *Toxoplasma* aspartyl protease (ASP)5, a Golgi-resident protease that is phylogenetically related to *Plasmodium* Plasmepsin V, mediates the export of GRAs to the host cytosol and can influence the localization of several GRAs to the PVM (26–28). To investigate whether GRAs that localize at the PVM or GRAs that are exported to the host cytosol activate the NLRP1 inflammasome, cell viability of Lewis rat macrophages infected with Δ*asp5* parasites was measured (**Figure 2A**). Compared to wild-type (WT) parasite infection, Δ*asp5* parasites induced less macrophage cell death, and Δ*asp5* parasites complemented with a Ty-tagged copy of *ASP5* regained the ability to induce cell death (**Figure 2A, right panel**). MYR1, a putative *Toxoplasma* PVM translocon, mediates the export of GRAs, including GRA16 and GRA24, into the host cytosol (29). Δ*myr1* parasites (**Figure S2A and C**) induced similar levels of Lewis rat macrophage cell death compared to WT parasites (**Figure 2B**). Taken together, *Toxoplasma*-induced Lewis rat macrophage cell death is ASP5-but not MYR1-dependent suggesting that GRAs that localize to the PVM, but not GRAs exported to the host cytosol, are likely mediators of Lewis macrophage cell death.

**Figure 2.**
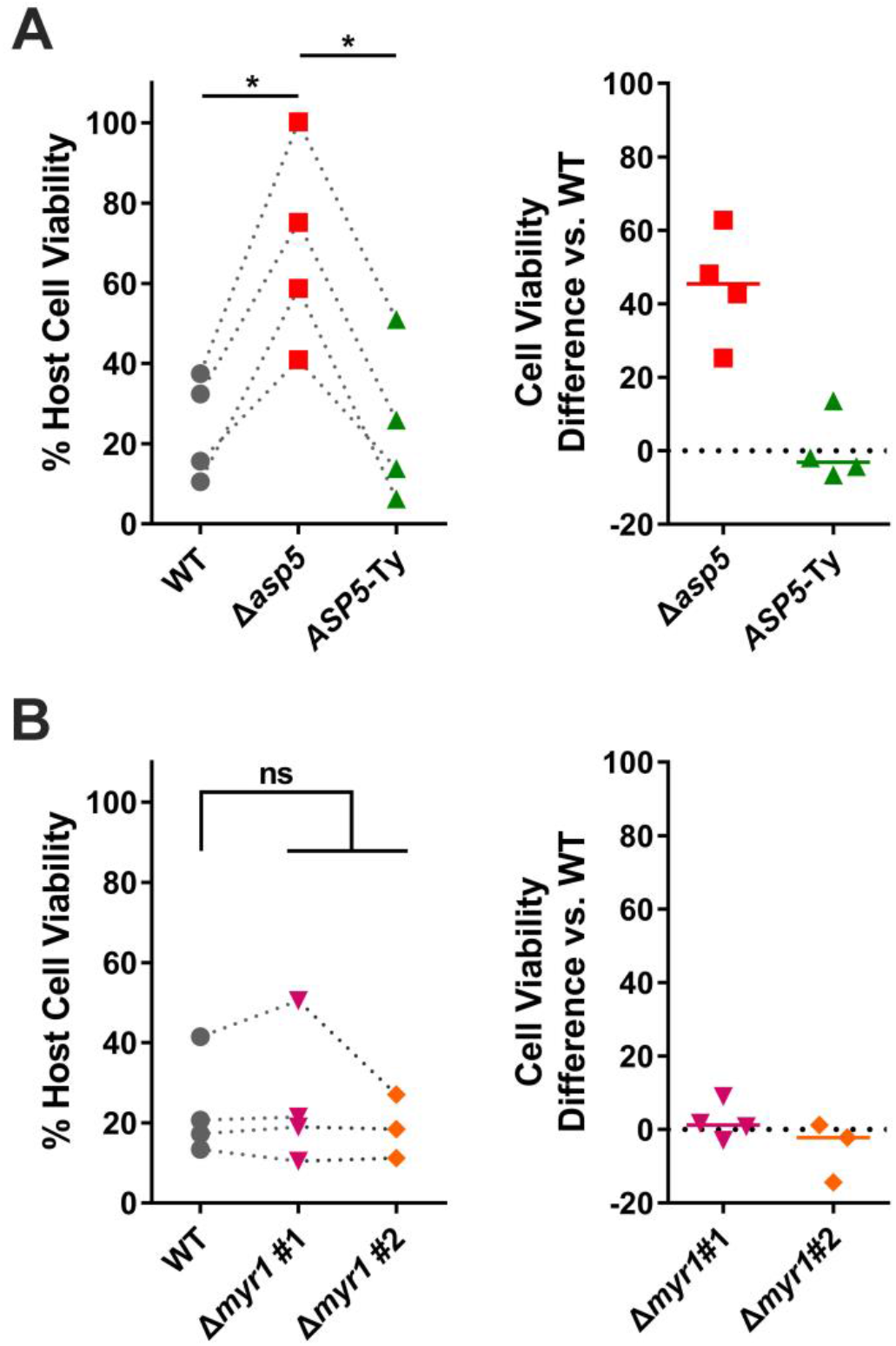
Toxoplasma-induced Lewis rat macrophage cell death is ASP5-but not MYR1-dependent. (**A**) Lewis rat BMDMs were infected with WT parasites, ASP5 knockout parasites (Δ*asp5*) or ASP5 knockout parasites complemented with a Ty-tagged copy of *ASP5* (*ASP5*-Ty) (MOI = 1) for 24 hours. Macrophage viability was measured via MTS assay. Data are displayed as the paired scatterplots (left, *n* = 4; \**p* < 0.05; student’s t-test). The right scatterplots are showing the cell viability difference between indicated strains with WT parasites in each paired experiment. Horizontal bars represent the median cell viability difference. (**B**) Lewis rat BMDMs were infected with WT parasites or two independent clones of MYR1 knockout parasites (Δ*myr1* #1 and Δ*myr1* #2) (MOI = 1) for 24 hours. Macrophage viability was measured via MTS assay. Data are displayed as the paired scatterplots (left, *n* = 4 for WT and Δ*myr1* #1, *n* = 3 for Δ*myr1* #2; ns, not significant; student’s t-test). The right scatterplots show the cell viability difference between Δ*myr1* parasites and WT parasites in each paired experiment. Horizontal bars represent the median cell viability difference.

### Isolation of *Toxoplasma* mutants that do not induce Lewis macrophage cell death

Although GRAs that localize to the PVM are likely involved in NLRP1 inflammasome activation, the effector(s) are still unknown. To identify *Toxoplasma* gene product(s) required for NLRP1 inflammasome activation, we designed a chemical mutagenesis screen to isolate mutants that fail to induce Lewis rat macrophage cell death (**Figure 3A**). Type I (RH) parasites were mutagenized by *N*-ethyl-*N*-nitrosourea (ENU) or ethyl methanesulfonate (EMS), respectively. The populations of chemically mutagenized parasites were used to infect Lewis rat macrophages at a multiplicity of infection (MOI) of 0.2-0.3. *Toxoplasma*-induced macrophage cell death is a dominant trait, reinvasion of parasites into rare cells containing *Toxoplasma* mutants that do not activate the NLRP1 inflammasome would therefore still lead to macrophage cell death. Therefore, to inhibit reinvasion, extracellular parasites were washed from cells after 2 hours of infection and the media was replaced with fresh media that contained the glycosaminoglycan, dextran sulfate (DS), a glycan competitor that prevents host cell invasion by extracellular parasites (30). Parasites that retain the ability to induce macrophage cell death are released from the lysed cell into the supernatant, where the parasite is coated with DS, blocking re-invasion into a new host cell. Mutated parasites unable to induce macrophage cell death are able to replicate within the surviving macrophage. After 24 hours of infection, surviving cells were washed, thereby removing the extracellular parasites capable of inducing macrophage cell death from the population. The surviving macrophages were then added to a monolayer of human foreskin fibroblasts (HFFs) so the parasites within the macrophages could continue to replicate until their natural egress from the macrophages.

**Figure 3.**
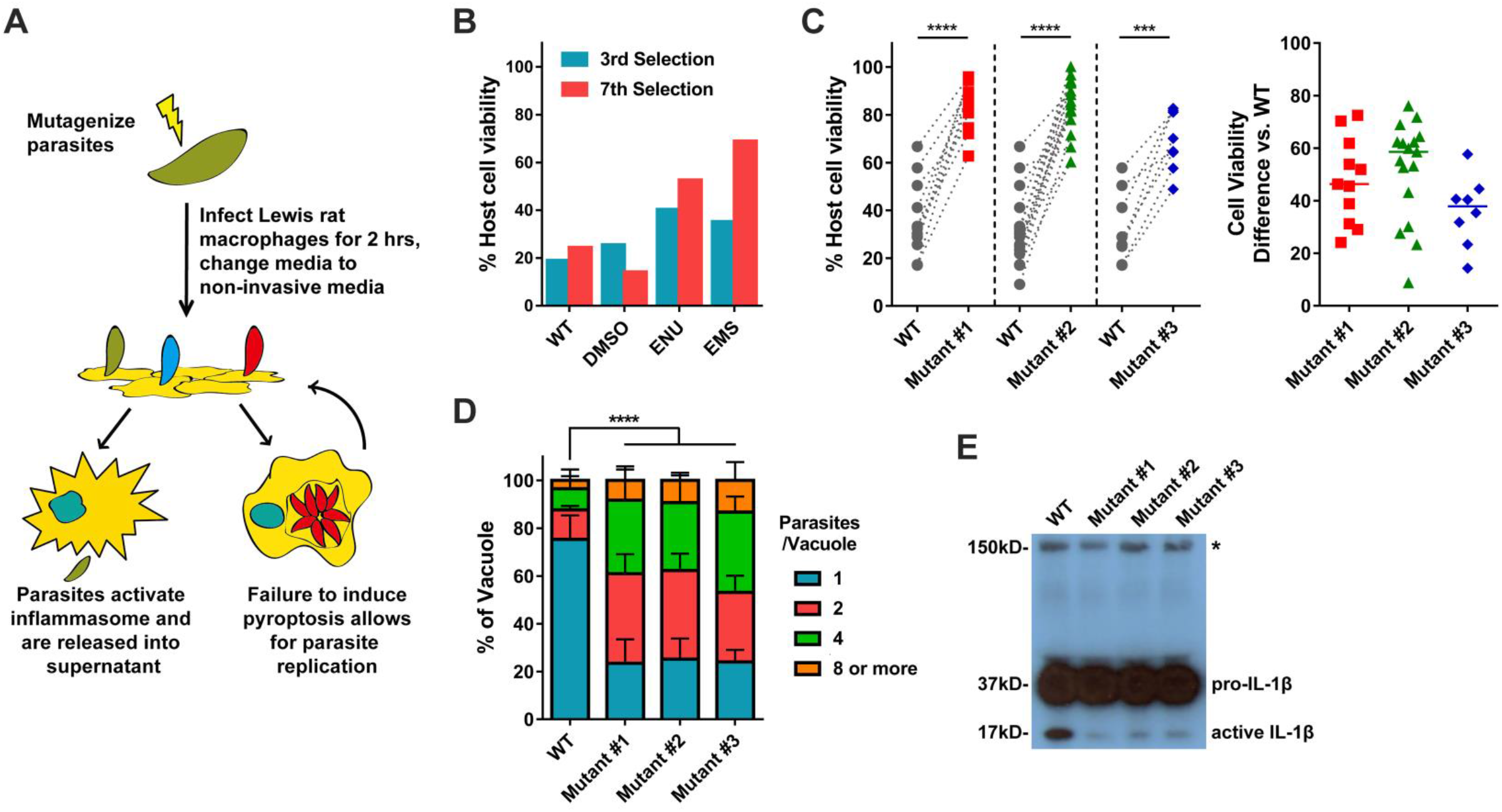
Isolation of *Toxoplasma* mutants that do not induce Lewis rat macrophage cell death. (**A**) Schematic of mutagenesis screen. DS is Dextran Sulfate, BMDMs is Bone marrow-derived macrophages. (**B**) Lewis rat BMDMs were infected with indicated mutagenized parasites (MOI = 1) for 24 hours. Macrophage viability was measured via MTS assay. Data are displayed as the column (*n* = 1). (**C**) Lewis rat BMDMs were infected with WT parasites or independent mutant strains isolated from the pool of mutagenized parasites (Mutant clone #1, #2 and #3) (MOI = 1) for 24 hours. Macrophage viability was measured via MTS assay. Data are displayed as the paired scatterplots (left, *n* ≥ 8 for WT, *n* = 11 for mutant #1, *n* = 17 for mutant #2, *n* = 8 for mutant #3; \*\*\**p* < 0.001, \*\*\*\**p* < 0.0001; student’s t-test). The right scatterplots are showing the cell viability difference between indicated mutant strains and WT parasites in each paired experiment. Horizontal bars represent the median cell viability difference. (**D**) Lewis rat BMDMs were infected with the strains used in (**C**) (MOI = 0.5) for 24 hours. Number of parasites per vacuole was quantified by microscopy. Between 100-120 vacuoles were counted per experiment. Data are displayed as the average values (*n* = 4; error bars, +SD; \*\*\*\**p* < 0.0001; two-way ANOVA comparing mutants to WT). (**E**) Western blot probing for IL-1β on concentrated (20x) supernatants of LPS-primed (100 ng/ml, 2 hours) Lewis rat BMDMs infected with the strains used in (**C**) (MOI = 1) for 24 hours. Image is representative of two experiments, pro-IL-1β is 37 kD, active IL-1β is 17 kD, aspecific band is represented by asterisk and indicates similar loading of samples.

After seven rounds of selection, a distinct phenotype (the cell viability of Lewis rat macrophages upon *Toxoplasma* infection is more than 50%) began to emerge in two independent populations of mutagenized parasites compared to WT and dimethyl sulfoxide (DMSO)-treated parasites (**Figure 3B**). After a further two rounds of selection, single parasites were cloned from the populations and individual clones were tested for their inability to induce Lewis macrophage cell death. Three independent mutant clones induced significantly less Lewis rat macrophage cell death (**Figure 3C**). Macrophage survival was linked to the ability of the parasite to replicate within the macrophage. As expected, 75% of the surviving macrophages infected with WT parasites contained only single parasites while only 25% of cells infected with the mutants contained single parasites (**Figure 3D**). Inflammasome activation is also characterized by active IL-1β secretion. We found a strong decrease in the amount of cleaved, active IL-1β (17 kD) secreted from macrophages infected with each of the mutant strains, compared to WT (**Figure 3E**). Thus, the forward genetic selection strategy was successful in yielding *Toxoplasma* mutants deficient in the activation of the inflammasome in Lewis rat macrophages.

### Identification of single nucleotide variations in the mutants

To identify the genes mutated in each clone, we performed whole genome sequencing of each mutant. Sequence comparisons relative to the parental strain revealed 16, 11 and 12 non-synonymous mutations in mutant #1, mutant #2 and mutant #3, respectively (**Table 1**). The three mutants did not have any mutated genes in common. To identify the causative mutations in these mutants, we established a set of criteria to narrow the list of possible genes. The inflammasomes are expressed and assembled within the cytoplasm of host cells. We therefore chose to focus on *Toxoplasma* genes whose protein products contain predicted signal peptides. Additionally, we previously tested a large number of different *Toxoplasma* strains for their ability to activate the inflammasome and all strains tested were able to induce pyroptosis (5). We therefore focused on genes that were expressed (FPKM>10) across all strains based on our published RNAseq dataset for these strains (31). Using these criteria, we narrowed the list of candidate genes in these mutants to seven genes (**Figure 4A**).

**Figure 4.**
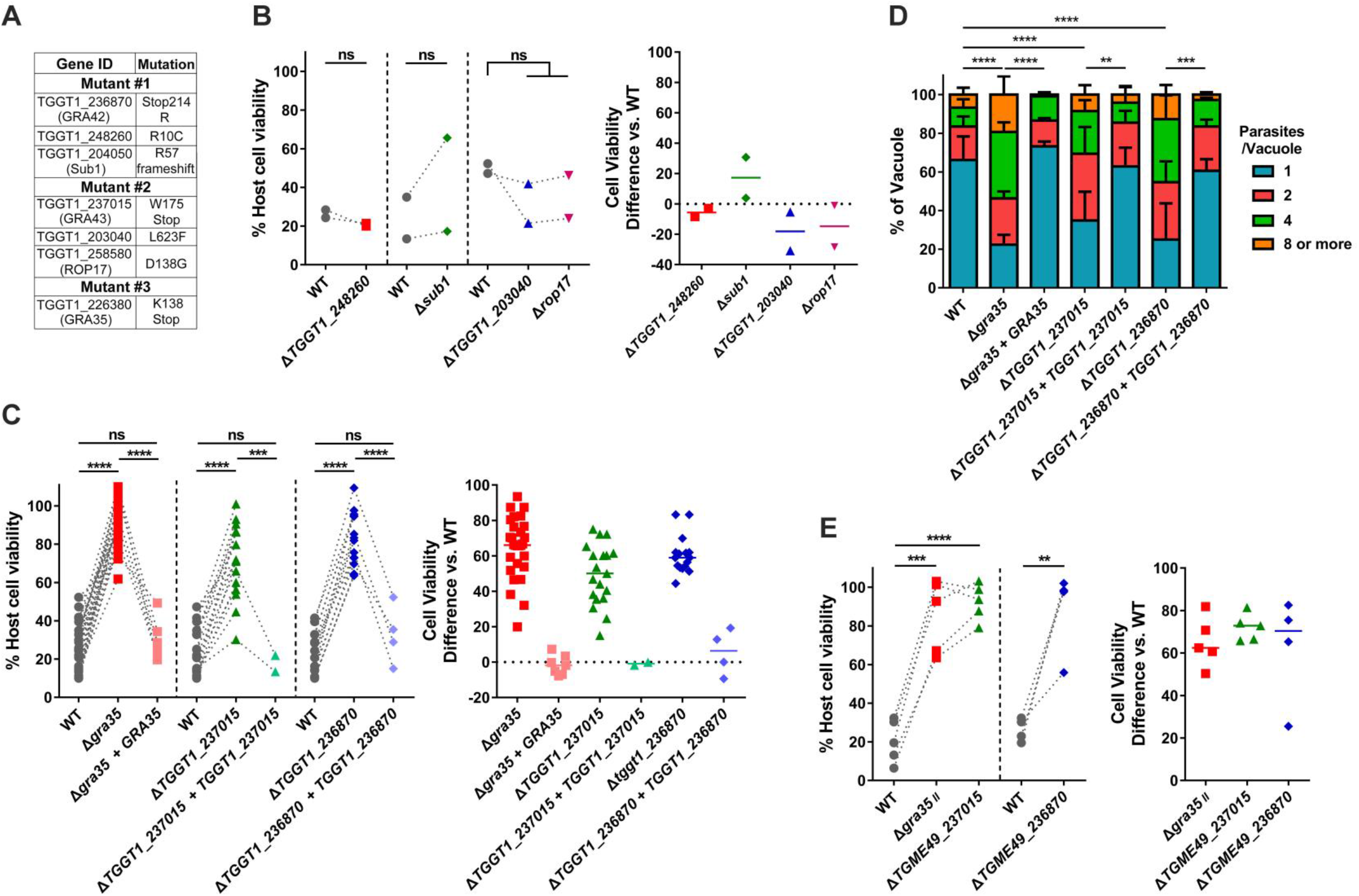
Three genes are individually required to induce cell death in Lewis rat BMDMs. (**A**) List of genes containing non-synonymous polymorphisms that fulfill candidate gene criteria in isolated mutants. (**B**) Lewis rat BMDMs were infected with WT parasites or the parasites in which *TGGT1_248260, SUB1, TGGT1_203040* or *ROP17* was knocked out (Δ*TGGT1_248260*, Δ*sub1*, Δ*TGGT1_203040* or Δ*rop17*) (MOI = 1) for 24 hours. Macrophage viability was measured via MTS assay. Data are displayed on the left as the paired scatterplots (Left, *n* = 2; ns, not significant; student’s t-test). The right scatterplots are showing the cell viability difference between indicated knockout strains and WT parasites in each paired experiment. Horizontal bars represent the median cell viability difference. (**C**) Cell viability as assessed by MTS assay of Lewis rat BMDMs infected with WT parasites, or parasites in which *GRA35, TGGT1_237015* or *TGGT1_236870* was knocked out (Δ*gra35*, Δ*TGGT1_237015* or Δ*TGGT1_236870*) or knockout parasites complemented with WT alleles of *GRA35, TGGT1_2370l5* or *TGGT1_236870* (Δ*gra35* + *GRA35*, Δ*TGGT1_237015 + TGGT1_237015* or Δ*TGGT1_236870 + TGGT1_236870*) (MOI = 1) for 24 hours. Data are displayed on the left as the paired scatterplots (left, *n* ≥ 16 for WT, *n* = 28 for Δ*gra35, n* = 7 for Δ*gra35* + *GRA35, n* = 19 for Δ*TGGT1_237015, n* = 2 for Δ*TGGT1_237015 + TGGT1_237015, n* =16 for Δ*TGGT1_236870, n* = 4 for Δ*TGGT1_236870 + TGGT1_236870*; \*\*\**p* < 0.001, \*\*\*\**p* < 0.0001, ns, not significant; student’s t-test). The right scatterplots are showing the cell viability difference between indicated strains with WT parasites in each paired experiment. Horizontal bars represent the median cell viability difference. (**D**) Number of parasites per vacuole were measured in Lewis rat BMDMs infected with the strains used in (**C**) (MOI = 0.5) at 24 hours post-infection. Between 100-120 vacuoles were counted per experiment. Data are displayed as the average values (*n* = 5 for WT and Δ*TGGT1_237015, n* = 4 for Δ*gra35, n* = 3 for Δ*TGGT1_236870, n* = 2 for all the complementation strains; error bars, +SD; \*\**p* < 0.01, \*\*\**p* < 0.001, \*\*\*\**p* < 0.0001; two-way ANOVA multiple comparisons). (**E**) Lewis rat BMDMs were infected with type II WT parasites or type II parasites in which *GRA35, TGME49_237015* or *TGME49 236870* was knocked out (Δ*gra35_II_*, Δ*TGME49_237015* or Δ*TGME49 236870*) (MOI = 1) for 24 hours. Macrophage viability was measured via MTS assay. Data are displayed as the paired scatterplots (left, *n* ≥ 4 for WT, *n* = 5 for Δ*gra35_II_* and Δ*TGME49_237015, n* = 4 for Δ*TGME49 236870; **p* < 0.01, \*\*\**p* < 0.001, \*\*\*\**p* < 0.0001; student’s t-test). The right scatterplots are showing the cell viability difference between indicated knockout strains with WT parasites in each paired experiment. Horizontal bars represent the median cell viability difference.

**Table 1.** List of all identified non-synonymous mutations. “Ref” is reference nucleotide(s) in WT strain (GT1 v9.0). “Sub” is nucleotide variant(s). “Mut” is mutant clone number.

| Chromosome | Position | Ref | Sub | Codon Change | AA Change | Gene | Mut No. |
| --- | --- | --- | --- | --- | --- | --- | --- |
| TGGT1_chrXII | 3698939 | C | T | Cgt/Tgt | R/C | TGGT1_248260 | 1 |
| TGGT1_chrXI | 4323464 | A | G | cTc/cCc | L/P | TGGT1_314875 | 1 |
| TGGT1_chrX | 5454719 | A | T | Tga/Aga | */R | TGGT1_236870 | 1 |
| TGGT1_chrVIII | 3546892 | A | G | Aca/Gca | T/A | TGGT1_273510 | 1 |
| TGGT1_chrVIIb | 258249 | C | G | Ccg/Gcg | P/A | TGGT1_263360 | 1 |
| TGGT1_chrVIIb | 1300287 | A | G | tTc/tCc | F/S | TGGT1_262825 | 1 |
| TGGT1_chrVIIb | 4053654 | G | C | Ccg/Gcg | P/A | TGGT1_257500 | 1 |
| TGGT1_chrVIIa | 683027 | A | G | Ttc/Ctc | F/L | TGGT1_206550 | 1 |
| TGGT1_chrVIIa | 1666878 | A | G | tTg/tCg | L/S | TGGT1_204310 | 1 |
| TGGT1_chrV | 2683109 | A | C | Ttg/Gtg | L/V | TGGT1_284040 | 1 |
| TGGT1_chrIX | 1745808 | G | A | cCc/cTc | P/L | TGGT1_264890 | 1 |
| TGGT1_chrIX | 3803976 | T | C | Tct/Cct | S/P | TGGT1_290960 | 1 |
| TGGT1_chrIII | 527809 | A | T | aaA/aaT | K/N | TGGT1_252395 | 1 |
| TGGT1_chrIII | 1241431 | C | T | Gac/Aac | D/N | TGGT1_253870 | 1 |
| TGGT1_chrIb | 814454 | A | T | cTg/cAg | L/Q | TGGT1_208580 | 1 |
| TGGT1_chrVIIa | 2153702 | GGA | GA | gag/aga | E/R | TGGT1_204050 | 1 |
| TGGT1_chrIV | 2235861 | G | A | cGa/cAa | R/Q | TGGT1_301250 | 2 |
| TGGT1_chrVIIa | 2964132 | C | G | ttG/ttC | L/F | TGGT1_203040 | 2 |
| TGGT1_chrVI | 290424 | T | C | Aaa/Gaa | K/E | TGGT1_239130 | 2 |
| TGGT1_chrVI | 3356628 | G | C | Gga/Cga | G/R | TGGT1_243635 | 2 |
| TGGT1_chrV | 121175 | A | G | aTc/aCc | I/T | TGGT1_220175 | 2 |
| TGGT1_chrXII | 5959927 | A | C | cAt/cCt | H/P | TGGT1_278518 | 2 |
| TGGT1_chrVIII | 2061823 | T | C | Agt/Ggt | S/G | TGGT1_231410 | 2 |
| TGGT1_chrVIIb | 730342 | C | T | Cgt/Tgt | R/C | TGGT1_264140 | 2 |
| TGGT1_chrVIIb | 2573674 | G | A | cCa/cTa | P/L | TGGT1_260450 | 2 |
| TGGT1_chrVIIb | 3451345 | A | G | gAt/gGt | D/G | TGGT1_258580 | 2 |
| TGGT1_chrX | 5567109 | C | T | tGg/tAg | W/* | TGGT1_237015 | 2 |
| TGGT1_chrIX | 2023395 | T | C | gAa/gGa | E/G | TGGT1_264472 | 3 |
| TGGT1_chrV | 1043175 | T | C | Acg/Gcg | T/A | TGGT1_213610 | 3 |
| TGGT1_chrVI | 1514625 | A | T | gAt/gTt | D/V | TGGT1_240960 | 3 |
| TGGT1_chrVI | 674303 | C | T | aGa/aAa | R/K | TGGT1_239700 | 3 |
| TGGT1_chrVIIa | 1197377 | C | G | Gcc/Ccc | A/P | TGGT1_205160 | 3 |
| TGGT1_chrVIII | 2566377 | G | T | gaG/gaT | E/D | TGGT1_233120 | 3 |
| TGGT1_chrX | 1583637 | A | T | Aaa/Taa | K/* | TGGT1_226380 | 3 |
| TGGT1_chrX | 3027043 | T | A | Agc/Tgc | S/C | TGGT1_224280 | 3 |
| TGGT1_chrX | 401396 | T | C | Agt/Ggt | S/G | TGGT1_228210 | 3 |
| TGGT1_chrXI | 2517037 | A | G | cAc/cGc | H/R | TGGT1_312140 | 3 |
| TGGT1_chrXII | 1102624 | T | C | Aag/Gag | K/E | TGGT1_219070 | 3 |
| TGGT1_chrXII | 6691803 | T | C | aAg/aGg | K/R | TGGT1_277030 | 3 |

To determine which of these genes are involved in Lewis rat NLRP1 inflammasome activation, we individually disrupted each candidate gene in the RH background (**Figure S2A and C**) and tested the resulting strains for their inability to induce macrophage cell death. Parasites in which we knocked out *TGGT1_248260, SUB1, TGGT1_203040*, or *ROP17* induced similar Lewis rat macrophage cell death compared to WT parasites (**Figure 4B**). Mutant #3 has only one candidate gene, *TGGT1_226380*, encoding GRA35 (32). A mutation in this gene resulted in an early stop codon (**Figure 4A and Figure S3A**). In mutant #2, a mutation in *TGGT1_237015* also resulted in an early stop codon (**Figure 4A and Figure S3A**). In mutant #1 a mutation in the stop codon of *TGGT1_236870* converted this stop codon into an Arginine (R), which resulted in an extended gene product (**Figure 4A and Figure S3A**). Lewis rat macrophages infected with parasites that contained individual disruptions in *GRA35, TGGT1_237015* or *TGGT1_236870* showed significantly less cell death compared to macrophages infected with WT parasites (**Figure 4C**). Complementation of knockout strains with WT alleles of *GRA35, TGGT1_237015* and *TGGT1_236870* restored their ability to induce Lewis rat macrophage cell death (**Figure 4C**). The replication of Δ*gra35*, Δ*TGGT1_237015* and Δ*TGGT1_236870* parasites in infected Lewis rat macrophages was significantly enhanced compared to WT parasites and complemented parasites 24 hours after infection (**Figure 4D**). Similarly, type II (ME49) parasites in which *GRA35, TGME49_237015* or *TGME49_236870* were disrupted (**Figure S2C**) induced less Lewis macrophage cell death compared to macrophages infected with WT parasites (**Figure 4E**). We also sequenced these 3 genes in other independent mutants. Another mutation (Y121 mutant to stop codon) in *GRA35* was also found in one of these mutant clones (named mutant #4), which failed to induce Lewis rat macrophage cell death (**Figure S3A and Figure S4**). These results indicated that the gene products of *GRA35, TGGT1_237015* and *TGGT1_236870* mediate *Toxoplasma*-induced Lewis rat macrophage cell death.

### *TGGT1_236870* and *TGGT1_237015* encode for novel PV-localized dense granule proteins

GRA35 was identified as a novel PV-localized dense granule protein by Bio-ID using other GRAs as baits (32) but there are no reports on the gene products encoded by *TGGT1_237015* and *TGGT1_236870. GRA35, TGGT1_237015* and *TGGT1_236870* are small one exon genes that are expressed in all *Toxoplasma* life stages except in the sexual stages inside the cat (www.toxodb.org). The predicted protein products of these genes lack predicted functional domains except for the C-terminal coiled-coil domain of GRA35 (**Figure S3A**). The resulting proteins each have a signal peptide, one predicted transmembrane (TM) domain and are generally predicted to be very alpha helical except the gene product of *TGGT1_236870* (**Figure S3A**). No *Toxoplasma* export element (TEXEL, RRLxx) motif (26) is present in the amino acid sequence of GRA35, TGGT1_237015 and TGGT1_236870. Although these three genes are quite conserved among different *Toxoplasma* strains, the rates of non-synonymous/synonymous (NS/S) polymorphisms between 64 different strains are higher at the C-terminus (starting after the TM domain) of each gene product (**Figure S3B to D**). BLAST analysis of the entire protein sequence revealed no predicted function of these three genes. Orthologs of *GRA35, TGGT1_237015* and *TGGT1_236870* were identified in other tissue cyst-forming coccidia, *Hammondia hammondi, Neospora caninum* and *Besnoitia besnoiti* (**Figure S5**). We also found that three *Toxoplasma* proteins, TGGT1_225160, GRA36 (TGGT1_213067) and TGGT1_257970, shared high amino acid similarity (>40%) with GRA35 (**Figure S5A**). Parasites deficient in *TGGT1_225160, GRA36* or *TGGT1_257970* still induced similar level of Lewis rat macrophage cell death compared to infection with WT parasites suggesting these proteins do not share the GRA35 function that mediates Lewis rat NLRP1 inflammasome activation (**Figure S6**).

To characterize GRA35, TGGT1_237015 and TGGT1_236870, we used complemented strains in which a C-terminally hemagglutinin (HA)-tagged version of each gene product is ectopically expressed in the knockout strains. The expression of each protein was confirmed by Western blot (**Figure 5A**). Both extracellular and intracellular parasites yielded a band migrating at identical size, suggesting GRA35, TGGT1_237015 and TGGT1_236870 did not undergo proteolytic modification in the process of secretion. The subcellular localization of each protein was observed in extracellular parasites. As previously reported, GRA35 is a dense granule protein that localized at punctuate structures which overlap with GRA7 while being excluded from rhoptries (**Figure 5B**). The gene products of *TGGT1_237015* and *TGGT1_236870* also showed co-localization with GRA7 but not ROP1 (**Figure 5B**). The three proteins were localized at the PVM and PV lumen in intracellular parasites (**Figure 7B, upper row**) suggesting they are indeed secreted *via* dense granules. We concluded from these data that TGGT1_236870 and TGGT1_237015 are novel dense granule proteins and therefore we named them GRA42 and GRA43, respectively.

**Figure 5.**
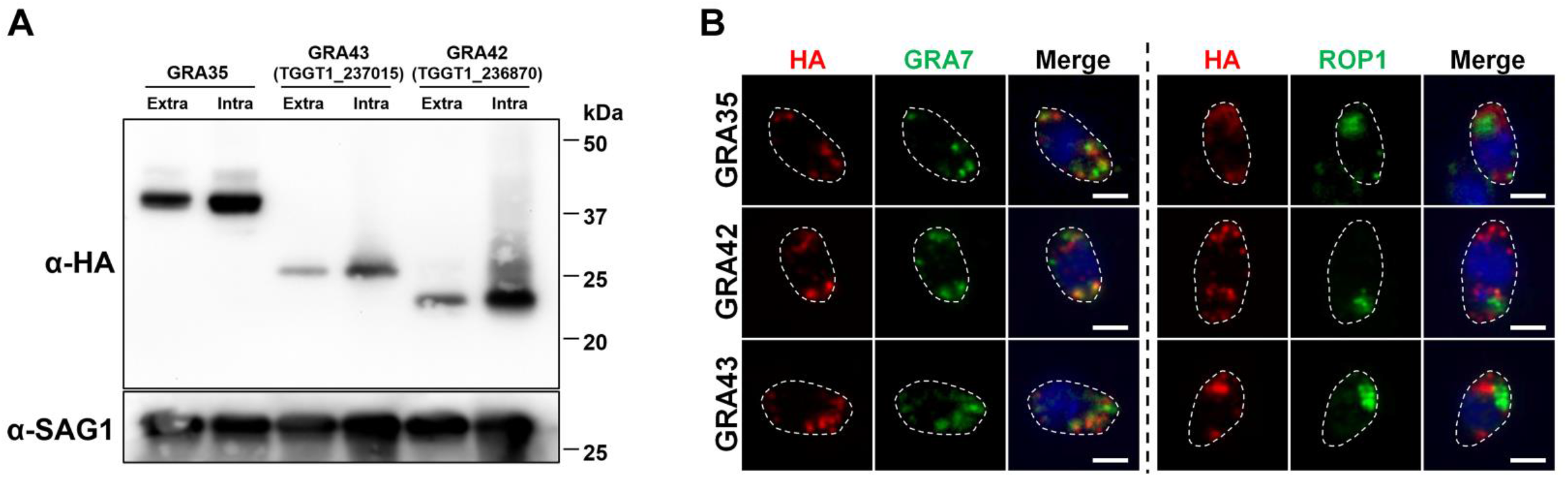
*TGGT1_236870* and *TGGT1_237015* encode for novel dense granule proteins, GRA42 and GRA43. (**A**) Strains individually knocked out in each gene were generated using CRISPR/Cas9 and complemented with an HA-tagged WT version of gene. HFFs were infected with HA-expressing parasites for 24 hours. Extracellular parasites were removed and washed with PBS prior to lysing (“Extra”). Remaining infected cells were lysed (“Intra”). SAG-1 is used as parasite loading control. Predicted sizes: GRA35, 40.3 kD; GRA42, 29.3 kD; GRA43, 23.8 kD. Image is representative of two independent experiments. (**B**) Extracellular parasites expressing HA-tagged GRA35, GRA42 or GRA43 were fixed, permeabilized, and subjected to Immunofluorescent assay with antibodies indicated. The images were taken at identical exposure times for each channel (scale bar = 2 μm). Image is representative of two independent experiments.

### Complementation of mutants with GRA35, GRA42 and GRA43 restores Lewis rat inflammasome activation

To confirm that the mutation in GRA35, GRA42 and GRA43 was indeed responsible for the failure to activate the inflammasome by our chemically mutagenized parasites, we expressed the WT allele of the gene in each mutant. Addition of WT version of *GRA35, GRA42* and *GRA43* to their respective mutant was sufficient to restore induction of Lewis rat macrophage cell death (**Figure 6A**). Similarly, macrophages infected with mutant strains expressing WT version of *GRA35, GRA42* or *GRA43* contained less replicating parasites compared to mutant-infected BMDMs (**Figure 6B**). We also observed an increase in the active IL-1β secreted from macrophages infected with the complemented strains compared to their mutant counterparts (**Figure 6C**). Overall these data indicate that GRA35, GRA42 and GRA43 are required for activation of the Lewis rat NLRP1 inflammasome by *Toxoplasma*.

**Figure 6.**
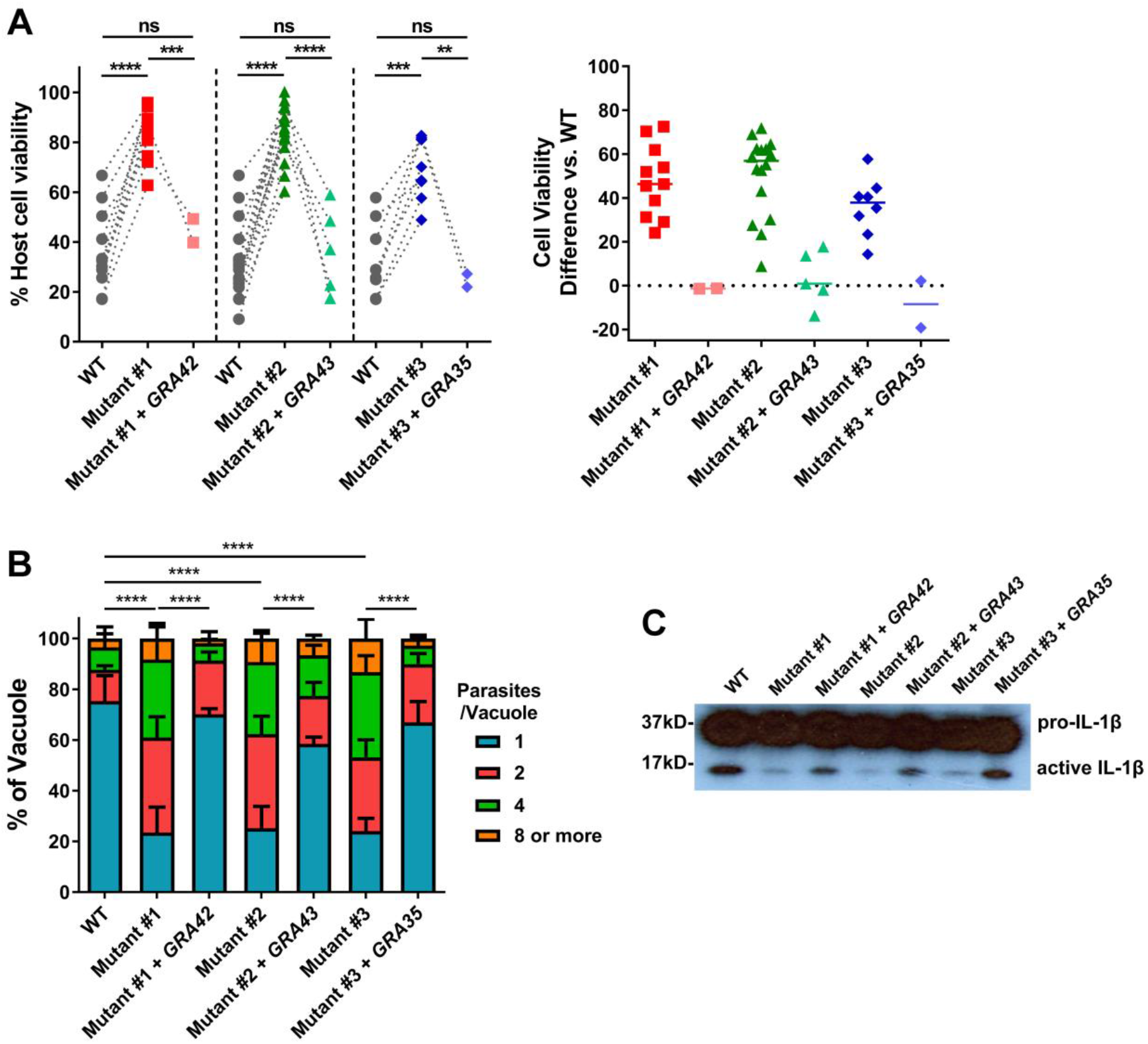
GRA35, GRA42 and GRA43 restore the mutant phenotype, and are required for inflammasome activation. (**A**) Lewis rat BMDMs were infected with WT parasites, independent mutant strains isolated from the pool of mutagenized parasites (Mutant clone #1, #2 and #3) or the mutant strains complemented with WT alleles of *GRA42, GRA43* or *GRA35* (Mutant #1 + *GRA42*, Mutant #2 + *GRA43*, Mutant #3 + *GRA35*) (MOI = 1) for 24 hours. Macrophage viability was measured via MTS assay. Data are displayed on the left as the paired scatterplots (left, *n* ≥ 8 for WT, *n* = 11 for mutant #1, *n* = 17 for mutant #2, *n* = 8 for mutant #3, *n* = 2 for mutant #1 + *GRA42* and mutant #3 + *GRA35, n* = 4 for mutant #2 + *GRA43; **p* < 0.01, \*\*\**p* < 0.001, \*\*\*\**p* < 0.0001; student’s t-test). The right scatterplots are showing the cell viability difference between indicated strains and WT parasites in each paired experiment. Horizontal bars represent the median cell viability difference. (**B**) Number of parasites per vacuole were measured in Lewis rat BMDMs infected with the strains used in (**A**) (MOI = 0.5) at 24 hours post-infection. Between 100-120 vacuoles were counted per experiment. Data are displayed as the average values (n = 4 for WT and mutant #1, #2 and #3, *n* = 2 for mutant #1 + *GRA42*, mutant #2 + *GRA43* and mutant #3 + *GRA35;* error bars, +SD; \*\*\*\**p* < 0.0001; two-way ANOVA multiple comparisons). (**C**) Western blot of IL-1β on concentrated supernatants (20x) BMDMs primed with LPS (100ng/ml, 2 hours) infected with the strains used in (**A**) (MOI =1, 24 hours). Image is representative of two independent experiments.

### GRA42 and GRA43 influence the PVM localization of GRA35, as well as other PVM-localized GRAs

Lewis rat macrophages infected with individual knockouts of *GRA35, GRA42* or *GRA43* showed a similar level of reduced cell death compared to macrophages infected with WT parasites (**Figure 4C, right panel**). It is therefore likely that these three GRAs function in the same pathway that activates the NLRP1 inflammasome. To confirm this, we generated double and triple *GRA35, GRA42* and *GRA43* knockout parasites (**Figure S2B and C**). Single-, double-or triple- *GRA35, GRA42* and *GRA43* knockout parasites induced similar levels of macrophage cell death (**Figure 7A**) indicating that these GRAs function in the same pathway. Possibly they form a protein complex that directly activates the inflammasome, or one of the GRAs activates the inflammasome and the other two are upstream in the pathway. To investigate this, we first determined the exact localization of GRA35, GRA42 and GRA43 in intracellular parasites. GRA35 localized at the PVM, while GRA42 and GRA43 were predominantly localized in the PV lumen (**Figure 7B, upper row**). We then determined the localization of GRA35, GRA42 and GRA43 in the different knockout parasites. In Δ*gra42* and Δ*gra43* parasites, GRA35 was mostly retained in the PV lumen and less of it was localized to the PVM, whereas the localization of GRA42 and GRA43 was unchanged regardless of the presence of GRA35, GRA42 or GRA43 (**Figure 7B, middle two row**). Previously, we found parasites deficient in ASP5 induced less Lewis rat macrophage cell death (**Figure 2A**). *ASP5* deletion also resulted in mis-localization of certain PVM-localized GRAs (26, 27). To understand whether ASP5 might influence Lewis rat macrophage cell death through these GRAs, the localization of GRA35, GRA42 and GRA43 was also observed in parasites lacking ASP5. In Δ*asp5* parasites GRA35 no longer localized to the PVM and was mostly present in the PV space (**Figure 7B, left bottom**). In contrast, ASP5 did not influence the localization of GRA42 and GRA43 (**Figure 7B, middle and right bottom**). Therefore, these results revealed that GRA42, GRA43, and ASP5 influence the PVM localization of GRA35. To understand whether GRA35 is the only GRA of which the localization is influenced by GRA42 and GRA43, we determined the localization of GRA17 and GRA23, which are also PVM-localized GRAs, in Δ*gra42* or Δ*gra43* parasites (**Figure 7C**). In WT parasites, these two GRAs were entirely localized at the PVM (**Figure 7C, top row**). In Δ*gra42* parasites, GRA17 and GRA23 were mis-localized to the PV space, although a small fraction localized to the PVM (**Figure 7C, middle row**). In Δ*gra43* parasites, these two GRAs were mostly absent at the PVM instead being retained in PV lumen (**Figure 7C, bottom row**). In contrast to GRA42 and GRA43, parasites deficient in GRA35 did not result in mis-localization of these two PVM GRAs. Note that only a small amount of GRA17 is required to mediate normal small molecule permeability and prevent enlarged vacuoles (33), possibly explaining why we failed to see the established Δ*gra17* ‘bubble vacuole’ phenotype in these vacuoles. Partial or no GRA17/GRA23 PVM staining was observed in more than 80% of the vacuoles of Δ*gra42* and Δ*gra43* parasites (**Figure 7D**). Therefore, GRA42 and GRA43 not only influence GRA35 localization at the PVM but also affect the localization of other PVM-associated GRAs.

**Figure 7.**
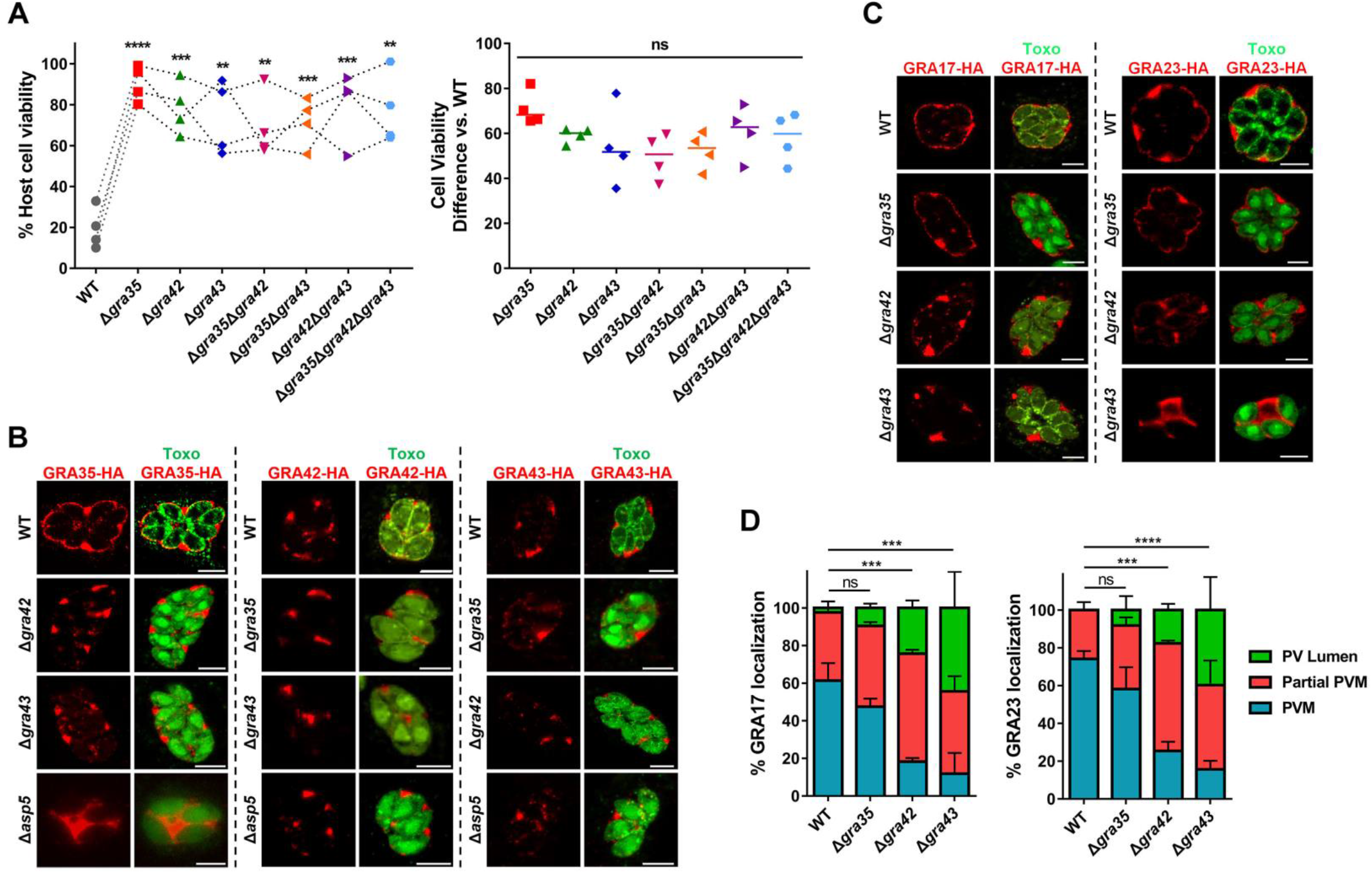
GRA42 and GRA43 influence the localization of GRA35, as well as GRA17, to the PVM. (**A**) Lewis rat BMDMs were infected with WT parasites, or parasites in which *GRA35, GRA42* or *GRA43* was knocked out (Δ*gra35*, Δ*gra42* or Δ*gra43*), or parasites containing a doubly knockout of *GRA35, GRA42* or *GRA43* (Δ*gra35*Δ*gra42*, Δ*gra35*Δ*gra43* or Δ*gra42*Δ*gra43*) or triple knockout parasites (Δ*gra35*Δ*gra42*Δ*gra43*) (MOI = 1) for 24 hours. Macrophage viability was measured via MTS assay. Data are displayed as the paired scatterplots (left, *n* = 4; all knockout strains vs. WT, \*\**p* < 0.01, \*\*\**p* < 0.001, \*\*\*\**p* < 0.0001; student’s t-test). The right scatterplots are showing the cell viability difference between indicated strains and WT parasites in each paired experiment. Horizontal bars represent the median cell viability difference (ns, not significant; one-way ANOVA with Kruskal-Wallis test). (**B**) HFFs were infected with WT parasites, parasites in which *GRA35, GRA42, GRA43* or *ASP5* was knocked out (Δ*gra35*, Δ*gra42*, Δ*gra43* or Δ*asp5*) and that transiently expressed GRA35-HA (left), GRA42-HA (middle) or GRA43-HA (right). The parasites were fixed and stained with antibodies against the HA epitope (red) and SAG1 (green). Transfected parasites were GFP positive. Images were taken at identical exposure times for each channel (scale bar = 5 μm). Image is representative of two independent experiments. (**C**) HFFs were infected with WT parasites or the parasites in which *GRA42* or *GRA43* was knocked out (Δ*gra42* or Δ*gra43*) and that transiently expressed GRA17-HA (left) or GRA23-HA (right), fixed and stained with antibodies against SAG1 (green) and the HA epitope (red). Transfected parasites were GFP positive. The images were taken at identical exposure times for each channel (scale bar = 5 μm). Image is representative of two independent experiments. (**D**) Localization of GRA17 or GRA23 (**C**) in at least 60 vacuoles containing 4 or more parasites was observed and scored as PVM localization, partial PVM localization or PV lumen localization. Data are displayed as the average values (*n* = 2; error bars, +SD; \*\*\**p* < 0.001, \*\*\*\**p* < 0.0001; two-way ANOVA comparing mutants to WT).

### No interaction between *Toxoplasma* GRA35 and Lewis rat NLRP1 in co-transfected HEK293T cells

GRA35 localized onto the PVM where its C-terminus possibly directly interacts with host cytosolic NLRP1. Because cell death occurs rapidly after parasite invasion (Cirelli et al., 2014), it is hard to detect the interaction between GRA35 and NLRP1 in parasite-infected macrophages. To investigate a direct interaction between Lewis rat NLRP1 and *Toxoplasma* GRA35, coimmunoprecipitation was performed in HEK293T cells transiently expressing FLAG-NLRP1 and GRA35-HA. The lysis of co-transfected cells was subjected to immunoprecipitation by using HA antibody and FLAG antibody. However, GRA35-HA was not detected in the FLAG-immunoprecipitated fraction, nor was FLAG-NLRP1 detected in the HA-immunoprecipitated fraction (**Figure 8**). Thus, the Lewis rat NLRP1 does not directly interact with *Toxoplasma* GRA35 in co-transfected HEK293T cells.

**Figure 8.**
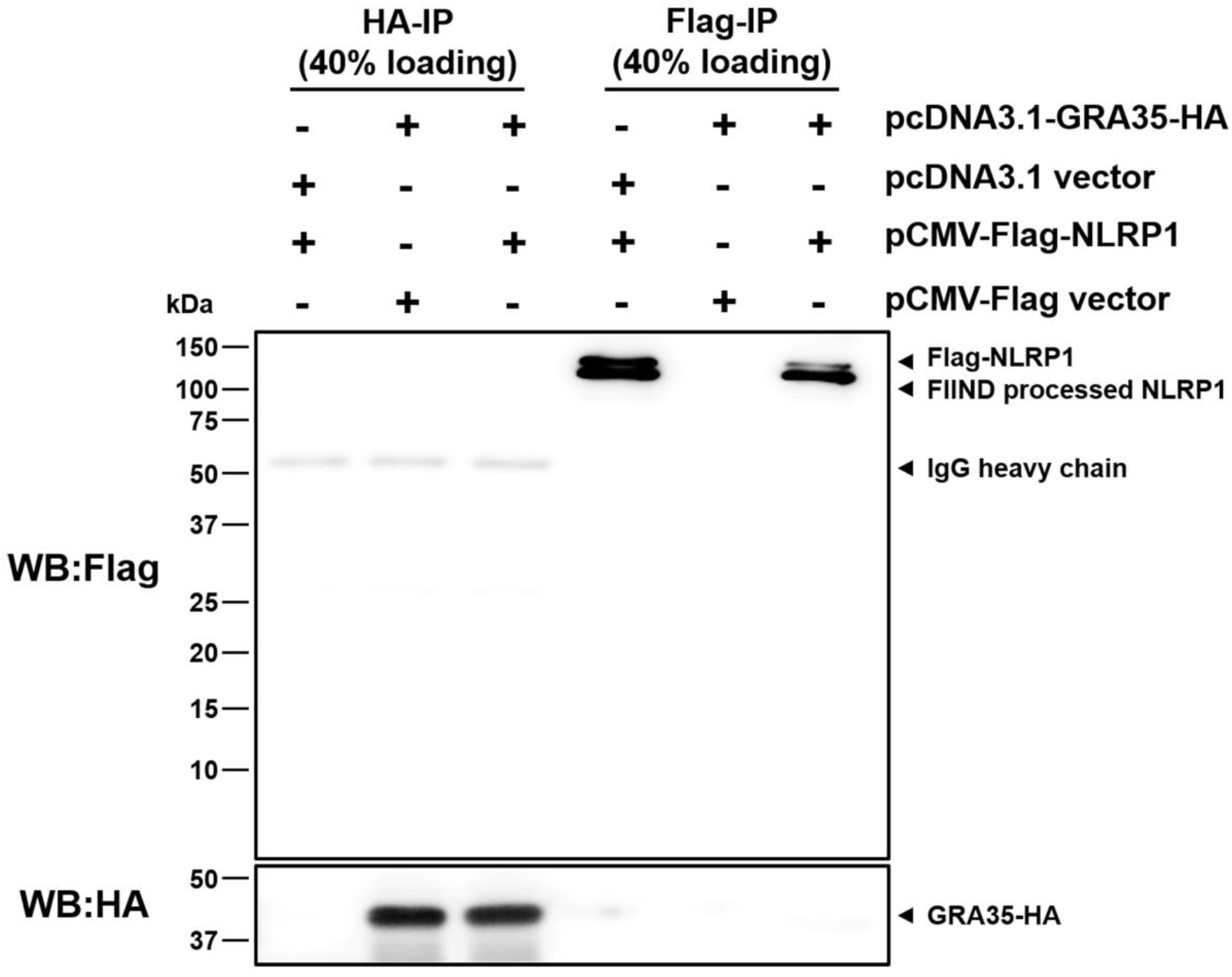
Lewis rat NLRP1 does not interact with *Toxoplasma* GRA35 in co-transfected HEK293T cells. HEK293T cells were co-transfected with pcDNA3.1-GRA35-HA and pCMV-FLAG-NLRP1 (expressing Lewis rat variant of Nlrp1) at the ratio of 1:1. 30 hours after transfection, cells were lysed in IP-lysis buffer (50 mM Tris pH 7.4, 150 mM NaCl, 0.5% Triton X-100) containing 1 × protease inhibitor and 1 mM PMSF. The indicated portion of cell lysates was incubated with protein G magnetic beads pre-bound with rat anti-HA or mouse anti-FLAG antibody at 4 °C for 1 hour with rotation. After washing with IP-lysis buffer, proteins bound to the beads were solubilized in SDS-loading buffer by boiling for 5 minutes, and examined by Western blot analysis using indicated antibody. Image is representative of two independent experiments with similar outcomes.

### *Toxoplasma* deficient in GRA35, GRA42 or GRA43 do not establish chronic infection in Lewis rats, but have reduced fitness in the rats that *Toxoplasma* does not activate the NLRP1 inflammasome

Since GRA35, GRA42 and GRA43 are required for activation of the NLRP1 inflammasome and parasite-induced pyroptosis in macrophages *in vitro*, we hypothesized that *Toxoplasma* strains deficient in these genes will fail to induce macrophage cell death *in vivo*, allowing the parasite to replicate and eventually disseminate to the brain leading to chronic infection. Removal of these genes does not lead to a general defect in parasite fitness in HFFs (34). We also found no significant difference in *in vitro* growth between WT parasites and Δ*gra35*, Δ*gra42* or Δ*gra43* parasites in rat fibroblasts (**Figure S7A**). Lewis rats were intraperitoneally infected with the type II ME49 strain expressing RFP or the *GRA35, GRA42* or *GRA43* knockout strains generated in this background. In addition, susceptible F344 rats, which encode an NLRP1 protein resistant to *Toxoplasma-mediated* inflammasome activation (2, 4), were used as a control. Compare to Lewis rat macrophages, F344 rat macrophages did not undergo rapid cell death after infection with WT or Δ*gra35*, Δ*gra42* or Δ*gra43* parasites (**Figure S7B**). During the course of infection, none of the rats lost weight or showed obvious clinical symptoms of toxoplasmosis (**data not shown**). After 2 months, the rats were sacrificed and the presence of cysts in the brains was determined. Brains of F344 rats infected with ME49-RFP parasites contained an average of 293 cysts whereas, as expected, no detectable cysts were found in the brains of Lewis rats. F344 rats infected with Δ*gra35*, Δ*gra42* or Δ*gra43* parasites contained reduced cyst numbers (73 cysts, 55 cysts and 0 cysts per brain of rats infected with Δ*gra35*, Δ*gra42* or Δ*gra43* parasites, respectively) (**Figure 9A**). This suggests that Δ*gra35*, Δ*gra42* and Δ*gra43* parasites determine *in vivo* fitness independent of their role in inflammasome activation. This was expected for Δ*gra42* and Δ*gra43* as these parasites have a defect in correct trafficking of GRAs to the PVM and some PVM GRAs, such as GRA17, determine parasite fitness. The absence of parasites in the brain of Δ*gra43* parasite-infected F344 rats was confirmed by diagnostic PCR based on the *Toxoplasma B1* gene (**Figure 9B**), which is a repetitive sequence in its genome (35). Reduced cyst number in F344 rats could be due to a defect of Δ*gra35*, Δ*gra42* or Δ*gra43* parasites in cyst formation. However, Δ*gra35*, Δ*gra42* or Δ*gra43* parasites formed normal *in vitro* cysts under alkaline stress induction condition (**Figure S7C**), suggesting these GRAs play no role in cyst formation. Lewis rats infected with Δ*gra35*, Δ*gra42* or Δ*gra43* parasites did not contain any brain cysts. Because the Δ*gra35*, Δ*gra42* and Δ*gra43* parasites determine fitness independent of their role in inflammasome activation we cannot make conclusions on the role of NLRP1 inflammasome activation in Lewis rat sterile immunity to *Toxoplasma*.

**Figure 9.**
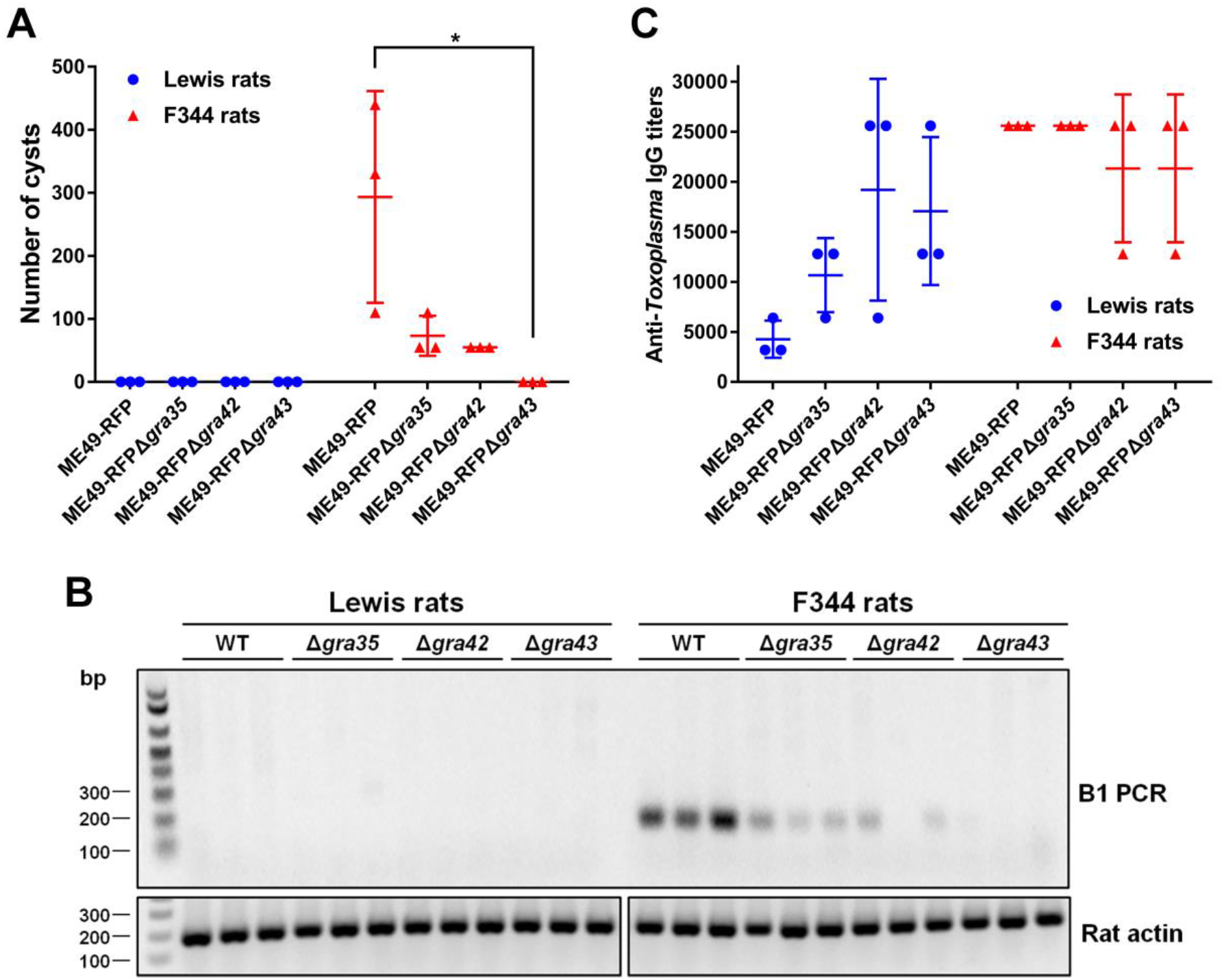
Parasites lacking GRA35, GRA42 and GRA43 do not establish a chronic infection in Lewis rats. (**A**) Number of brain cysts from each rat was determined by FITC-DBA staining at 60 days postinfection. Each plot represents number of brain cysts of individual rat (*n* = 3; \**p* < 0.05; one-way ANOVA with Kruskal-Wallis test). (**B**) The presence of *Toxoplasma* genomic DNA in the brain of infected rats was detected by diagnostic PCR targeting the multi-copy *B1* gene. As an internal control, rat actin was used to check the quality of isolated DNA. Image is representative of two independent experiments. (**C**) The rat serum was obtained at 60 days post-infection. The anti-*Toxoplasma* IgG titers were quantified by ELISA. Titers were defined as the dilution which gave an OD_405_ reading at least two-fold higher than the mean background in uninfected rat serum. Results are presented as mean values ± SD obtained from individual infected rats (*n* = 3).

Although Δ*gra35*, Δ*gra42* or Δ*gra43* seem to be generally much less virulent than WT in F344 rats we hypothesized that in Lewis rats their initial replication in macrophages might still allow them reach higher parasite numbers and disseminate compared to WT. Previously it was determined that higher parasite burdens in Lewis rats leads to higher anti-*Toxoplasma* antibody titers (2). We therefore compared the anti*-Toxoplasma* IgG titers in the sera obtained from all rats at 2 months post-infection (**Figure 9C**). Lewis rats infected with ME49-RFP parasites had lower anti*-Toxoplasma* IgG titers (1/3,200 - 1/6,400) compared to F344 rats (titers ≥ 1/25,600). Lewis rats infected with Δ*gra35*, Δ*gra42* or Δ*gra43* parasites had increased anti*-Toxoplasma* IgG titers (1/6,400 – 1/12,800, 1/6,400 – 1/25,600, 1/12,800 – 1/25,600, respectively) whereas titers were slightly decreased in F344 rats infected with Δ*gra42* or Δ*gra43* parasites (**Figure 9C**). The increased titers of Lewis rats infected with Δ*gra35*, Δ*gra42* or Δ*gra43*, compared to WT parasite infected rats, suggest that Δ*gra35*, Δ*gra42* or Δ*gra43* parasites bypassed the NLRP1 inflammasome barrier in macrophages allowing them to replicate and possibly disseminate. However, we were unable to observe detectable IL-1β level in the serum of parasite-infected Lewis rats and F344 rats regardless of parasite strain (**data not shown**). Taken together, even though GRA35, GRA42 and GRA43 are involved in activation of the Lewis rat NLRP1 inflammasome *in vitro, Toxoplasma* deficient in these genes still fail to develop cysts in the brain of Lewis rats likely because they are also required for *in vivo* fitness.

## Discussion

We and others previously showed that Lewis rat macrophage cell death upon *Toxoplasma* infection is determined by the NLRP1 inflammasome (4, 5). This study indicates that GRA35, GRA42 and GRA43 are parasite effectors that are involved in Lewis rat NLRP1 inflammasome activation. The fact that Δ*asp5* parasites, but not Δ*myr1* parasites, no longer induce *Toxoplasma*-induced Lewis rat macrophage cell death suggests that this cell death is mediated by PVM-localized GRAs. Several GRAs secreted onto the PVM have been identified as parasite effectors involved in host-parasite interactions including modulation of host signaling pathways, evasion of host immune responses, and nutrition acquisition (25). GRA6 locates at the PVM, where it selectively activates the host transcription factor nuclear factor of activated T cells 4 (NFAT4) via interaction with host Calcium modulating ligand (CAMLG) (36). GRA7 is a transmembrane protein that spans the PV and extends into the host cytosol, where it interacts with ROP complexes (37). GRA7 also binds directly to the oligomers of immunity-related GTPase Irga6 eventually leading to disassembly (37). GRA15 from type II *Toxoplasma*, another PVM-associated GRA, is involved in host NF-κB activation, which promotes the production of pro-inflammatory cytokines (38). Two additional dense granule proteins, GRA17 and GRA23, which are also located at the PV membrane, are responsible for small-molecule transport between the host cytosol and the vacuole lumen (33). Of the three GRAs we identified, only GRA35 localized at the PVM, while GRA42 and GRA43 are mainly localized inside the PV. This suggests that GRA35 might be the mediator of inflammasome activation as it has 1 TM domain and one part of GRA35 could face the host cytoplasm and possibly interact with the host cytosolic inflammasome. Since GRA35, GRA42 and GRA43 function in the same pathway that modulate inflammasome activation and GRA42 and GRA43 influence the PVM localization of GRA35 and other GRAs, it is likely that GRA42 and GRA43 function as protein chaperones that help GRAs localize to the PVM where GRA35 or another unknown GRA then activates the NLRP1 inflammasome either directly or indirectly.

Although our results indicate GRA35 could be the parasite effector that directly activates the Lewis rat NLRP1 inflammasome, the mechanism of activation is still unclear. Cleavage of NLRP1 is required for the activation of the inflammasome by Anthrax lethal toxin (20). A recent study demonstrated that proteolysis can act as a common activator of diverse NLRP1 variants from mice and humans (39). However, GRA35 does not have predicted protease domains and cleavage of NLRP1 was not found in GRA35 transfected-HEK293T cell line transiently expressing Lewis rat NLRP1, nor was a direct interaction between GRA35 and NLRP1 observed (**Figure 8**). *Toxoplasma* activation of mouse NLRP1 does not seem to involve cleavage of NLRP1 suggesting it might activate NLRP1 in mice through a novel mechanism (6). GRA35 has orthologues in *Hammondia, Neospora* and *Besnoitia. Neospora caninum* is able to induce cell death in Lewis rat macrophages (**Figure S8**), suggesting the function of GRA35 is conserved in cyst-forming coccidia.

The mutations of GRA35 in mutant #3 and mutant #4 are in the transmembrane domain, which results in GRA35 lacking its entire C-terminus containing two coiled-coil domains. Coiled-coil domains function in many biological processes, including protein-DNA binding and protein-protein interaction (40). However, no directly interaction between Lewis rat NLRP1 and *Toxoplasma* GRA35 was found in co-transfected HEK293T cells. Previously, we described that parasite infection of murine macrophage cell lines or human fibroblasts stably expressing Lewis rat NLRP1 does not trigger cell death (5). This suggests that murine macrophages and human fibroblasts lack a factor needed for activation of the Lewis rat NLRP1 inflammasome by *Toxoplasma*. Unfortunately, this also prevents us from using non-Lewis rat cell lines to determine if transfection of GRA35 is sufficient to activate the NLRP1 inflammasome. Overall our data are consistent with GRA35 interacting with a rat-specific factor that subsequently mediates the activation of the NLRP1 inflammasome. This pattern has been demonstrated for GRA6, whose C-terminus interacts with host cytosolic protein CAMLG, which leads to NFAT4 activation (36). It is also possible that GRA35 interacts with or modifies a Lewis rat-specific protein, which is sensed by NLRP1, similar to NLRC4 recognition of a NAIP5/NAIP6/flagellin complex (14, 41), or possibly inhibits the negatively regulation of NLRP1 by this rat factor. A further complication is that some inflammasomes do not directly interact with a PAMP but rather sense changes to the cellular milieu induced by infection. For example, NLRP3 senses diverse cellular signals, such as K^+^ efflux, Ca^2+^ signaling, reactive oxygen species (ROS), mitochondrial dysfunction, and lysosomal rupture, which are the triggers for NLRP3 inflammasome activation (42). It is therefore possible that NLRP1 does not directly interact with a *Toxoplasma* effector but rather detects changes in the cell induced by *Toxoplasma* infection. For instance, cytosolic ATP depletion is sensed by NLRP1b leading to inflammasome activation (43, 44). Another hypothesis is that GRA35 maybe function as a PVM platform that supports or modifies other parasite effectors that somehow activate the NLRP1 inflammasome. This model has been described for the ROP5/ROP18/ROP17/GRA7 complex, which locates at the PVM and prevents PVM rupture by preventing the accumulation of Immunity related GTPases (IRGs) (37, 45).

Although ASP5 influences GRA35 localization, there is no TEXEL motif present in GRA35 or in GRA42 and GRA43 suggesting that these three proteins are not direct substrates of ASP5. It is likely that another protein with a TEXEL motif mediates GRA35 localization to the PVM, or functions as a regulator of GRA42 and GRA43 function. Identification of this protein could help us gain a better understanding of how GRA35, GRA42 and GRA43 activate the Lewis rat NLRP1 inflammasome.

Unexpectedly, parasites lacking GRA35 were still unable to establish a chronic infection in Lewis rat. Because GRA42 and GRA43 are important for correct localization of other GRAs at the PVM (**Figure 7B and C**), some of which determine parasite fitness, it was expected that parasites lacking GRA42 or GRA43 would be less virulent *in vivo*, which makes it difficult to establish their role in NLRP1 activation on parasite *in vivo* fitness. Despite the failure in tissue cyst formation, the higher anti*-Toxoplasma* IgG titers in the serum of Lewis rats infected with Δ*gra35*, Δ*gra42* or Δ*gra43* parasites possibly indicates that the Lewis rat inflammasome was not activated during acute infection, allowing a limited proliferation of tachyzoites but that these parasites were eventually eliminated by other mechanisms. However, no parasites were detected in peritoneal organs (spleen and liver) or peritoneal cavity of Lewis rats and *Toxoplasma*-susceptible F344 rats by B1-PCR and *in vivo* imaging at 2 days post-infection (**data not shown**), suggesting the rat in general is resistant to the initial stage of infection. It remains unclear what mechanisms mediate parasite resistance in rats in which *Toxoplasma* does not activate the NLRP1 inflammasome (e.g. F344 rats). Because parasites lacking GRA35, GRA42 or GRA43 also had a defect in tissue cyst formation in susceptible F344 rats, which possess a Toxoplasma-resistant variant of *Nlrp1*, GRA35, GRA42 and GRA43 also have an inflammasome-independent role in the pathogenesis of the parasite *in vivo*.

Overall, the results presented here show that three dense granule proteins of *Toxoplasma gondii* are necessary for Lewis rat NLRP1 inflammasome activation. How these proteins function to activate the NLRP1 inflammasome is not yet known, but the data suggest a model where GRA42 and GRA43 mediate GRA35 localization to the PVM, where the GRA35 faces the host cytosol and mediates indirectly the activation of the NLRP1 inflammasome. Future experiments will be needed to determine the precise mechanism by which GRA35 mediates the activation of the NLRP1 inflammasome and GRA42 and GRA43 influence the localization of GRAs to the PVM.

## Materials and Methods

### Ethics statement

All animal experiments were performed in strict accordance with the recommendations in the Guide for the Care and Use of Laboratory Animals of the National Institutes of Health and the Animal Welfare Act, approved by the Institutional Animal Care and Use Committee at UC Davis (assurance number A-3433-01).

### Reagents and antibodies

ENU and EMS were purchased from Sigma-Aldrich. CellTiter 96 AQueous One Solution Cell Proliferation Assay was obtained from Promega. Dextran sulfate sodium salt was obtained from Santa Cruz Biotechnology. LPS was purchased from Calbiochem/EMD Biosciences. Caspase-1 inhibitor VX765 was purchased from Selleck chemicals. NLRP3 inflammasome inhibitor MCC950 was purchased from AdipoGen Life Sciences, Inc. Nigericin (Sodium Salt) was purchased from MilliporeSigma. Rabbit anti-IL-1β was purchased from Abcam. Rat anti-HA (3F10) antibody was obtained from Roche. Mouse anti-FLAG M2 antibody was purchased from Sigma-Aldrich. Secondary HRP-conjugated antibodies were purchased from Jackson ImmunoResearch. Alexa Fluor 448 and 594 secondary antibodies were obtained from Invitrogen.

### Rats and Parasites

Lewis (LEW/Crl; LEW) rats and F344 (F344/DuCrl; CDF) rats were purchased from Charles River Laboratories (Wilmington, MA) at 6-8 weeks old. Lewis rat bone marrow-derived macrophages (BMDMs) were prepared as previously described (5). *Toxoplasma gondii* tachyzoites from Type I (RH) expressing luciferase and GFP were used for mutagenesis. RH parasites without, luciferase and lacking the *HXGPRT* gene (RHΔhxgprt) were used for generating knockouts. RH parasites without, luciferase and lacking the *HXGPRT* gene and *Ku80* gene (*RHΔhxgprtΔku80*) were used as WT control of Δ*asp5* parasites. Type II (ME49) engineered to express RFP were a gift from Dr. Michael Grigg. RHΔ*sub1* was a kind gift from Dr. Vern Carruthers and generated as described (46). RHΔ*asp5* and RH-*A5P5*-Ty were kind gifts from Dr. Mohamed-Ali Hakimi and generated as described (28). All parasite strains were routinely passaged *in vitro* in monolayers of HFFs.

### Mutagenesis Screen

Intracellular RH parasites expressing GFP and Luciferase were treated with ENU (40 μM), EMS (100 μM) or DMSO for 4 hours. Parasites were washed three times with PBS, syringe lysed and allowed to infect fresh HFFs. For selection, Lewis BMDMs were infected with parasite populations (MOI = 0.2 – 0.3) for 2 hours. Non-invading parasites were removed by washing cells with PBS three times. Media was replaced with media containing 30 mg/ml dextran sulfate. At 24 hours post-infection, extracellular parasites were removed by washing cells with PBS three times. Cells were scraped into fresh media and overlaid onto fresh HFFs. After nine rounds selection, parasites were cloned via serial dilution. Parasite DNA was isolated using Qiagen DNeasy Blood & Tissue Kit according to manufacturer’s protocol. Illumina sequencing was performed on Illumina HiSeq 2000 or MiSeq. Reads were aligned using type I GT1 (v9.0) as reference genome.

### Generation of parasite strains

Individual knockout of candidate genes was performed using CRISPR-Cas9. Sequences targeting candidate genes were cloned into the pSS013-Cas9 vector (47). The sequences are available in **Supplementary Table 1**. To generate the *MYR1* knockout strain and knockout strains for the candidate hits from sequenced mutant clones, plasmids containing sgRNAs were co-transfected with XhoI (New England Biolabs)-linearized pTKOatt, which contains the *HXGPRT* selection cassette (38), into RHΔhxgprt parasites at a ratio 10:1 (sgRNAs: linearized pTKOatt plasmid). 24 hours post-transfection, populations were selected with mycophenolic acid (50 μg/ml) and xanthine (50 μg/ml) and cloned by limiting dilution (**Figure S2A**). Knockout was assessed by PCR (**Figure S2C**). Complemented strains were generated by cloning the gene with its putative promoter (~2000 bp upstream of start codon) with a C-terminal hemagglutinin (HA)-tag sequence into pENTR using TOPO cloning (Invitrogen) and then into pTKOatt using LR recombination (Invitrogen) (38). Prior to transfection, plasmids were linearized using a restriction enzyme with a unique restriction site. Parasites were co-transfected with the linearized complemented plasmid and a plasmid containing the dihydrofolate reductase (DHFR) resistance cassette at a ratio of 20:1. 24 hours post-transfection, populations were selected with pyrimethamine (1 μM) and cloned by limiting dilution. Presence of the tagged gene was determined by immunofluorescent assay (IFA) and Western blot. To generate the double and triple knockout strains, Δ*gra35* parasites were co-transfected with separate plasmids containing sgRNAs against GRA42 or GRA43 together with NotI (New England Biolabs)-linearized pLoxp-DHFR-mCherry (48), which also contains a pyrimethamine resistance cassette, at a ratio of 5:1 (**Figure S2B**). After two rounds of pyrimethamine selection and limiting dilution cloning, the double and triple knockout parasites were assessed by PCR and confirmed by sequencing in both loci. GRA42 and GRA43 double knockout strain was generated from Δ*gra42* parasites by using same strategy. To generate *TGGT1_225160, GRA36* or *TGGT1_257970* knockout strain, plasmids containing sgRNAs were co-transfected with NotI (New England Biolabs)-linearized pLoxp-DHFR-mCherry at a ratio of 5:1 (**Figure S5A**). After two rounds of pyrimethamine selection and limiting dilution cloning, the knockout parasites were assessed by PCR (**Figure S5B**) and confirmed by sequencing.

### Cell viability, parasite per vacuole counts, IL-1β measurement

Lewis rat BMDMs were stimulated with or without 50 μM of VX765 or 10 μM of MCC950 for 2 hours followed by parasite infection. F344 rat BMDMs were infected with parasites for 24 hours. Cell viability was measured by MTS as previously described (5). Parasites per vacuole counts were performed as previously described (5). In LPS-primed BMDMs, the culture supernatants were collected for IL-1β measurement by ELISA as previously described (5). IL-1β in infected cell culture supernatants was also concentrated using Amicon filters (3 kD molecular weight cutoff) (Millipore) and detected by Western blot.

### Coimmunoprecipitation

Plasmids expressing a C-terminal HA-tagged GRA35 without signal peptide (pcDNA3.1-GRA35-HA) and N-terminal FLAG-tagged Lewis rat variant of NLRP1 (pCMV-FLAG-NLRP1) were mixed at the ratio of 1:1 and transfected to HEK293T cells using X-tremeGENE 9 DNA Transfection Reagent (Roche) according to the manufacturer’s instructions. As controls, cells were also transfected with GRA35-HA + FLAG empty vector and pcDNA3.1 empty vector + FLAG-NLRP1 under the same conditions. After 30 hours transfection, cells were lysed in IP-lysis buffer (50 mM Tris pH7.4, 150 mM NaCl, 0.5% Triton X-100) containing 1 × protease inhibitor and 1 mM PMSF. The cell lysates were incubated with protein G magnetic beads pre-bound with rat anti-HA or mouse anti-FLAG antibody at 4 °C for 1 hour with rotation. After washing with IP-lysis buffer, proteins bound to the beads were solubilized in SDS-loading buffer by boiling for 5 minutes and examined by Western blot analysis. GRA35-HA was detected by rat anti-HA antibody, FLAG-NLRP1 was detected by mouse anti-FLAG antibody.

### Western blot

To detect activated IL-1β, concentrated culture supernatants were separated on 12% SDS-PAGE gels and transferred to PVDF membrane (Bio-Rad., USA). To detect HA-tagged-GRA35, GRA42 or GRA43 expression, cell lysates made from intracellular parasites and extracellular parasites were separated onto 12% SDS-PAGE gels and transferred to PVDF membrane. To detect interaction between GRA35-HA and FLAG-NLRP1, the coimmunoprecipitated samples were separated onto 12% SDS-PAGE gels and transferred to PVDF membrane. Western blot analysis was performed as previously described (38).

### Invasion assay

Lewis rat BMDMs were stimulated with or without 50 μM of VX765 or 10 μM of MCC950 for 2 hours followed by parasite infection. After 30 minutes infection, a red/green invasion assay was performed as previously described for indirect immunofluorescence (49).

### Immunofluorescent assay

Extracellular parasites released from syringe-lysed HFFs were loaded onto coverslips and fixed with 100% ice cold methanol for 5 minutes. Colocalization studies were performed with anti-GRA7 or anti-ROP1 and anti-HA antibodies. Alexa Fluor 488 and 594 secondary antibodies were used as previously described (38). To determine the localization of GRAs inside host cells, HFFs were infected with the different parasite strains for 24-30 hours, fixed with 3% formaldehyde for 20 minutes, permeabilized with 0.2% Triton X-100, followed by staining with rat anti-HA antibodies (1/500 dilution) or mouse monoclonal antibodies against *Toxoplasma* surface antigen (SAG1). Alexa Fluor 488 and 594 secondary antibodies were used as previously described (38).

### *In vitro* cyst induction

Parasites were propagated in HFFs on coverslips under bradyzoite-inducing conditions (RPMI 1640 medium supplemented with 50 mM HEPES and 1% fetal bovine serum, pH 8.2, ambient CO2) for 3 days. Cells were then fixed with 100% ice-cold methanol, permeabilized with 0.2% Triton X-100, and the cysts were stained by FITC-DBA (Vector Laboratories).

### *In vivo* infection, cyst counting, diagnostic PCR and serological detection

*Toxoplasma* tachyzoites were harvested from cell culture and released by passage through a 27-gauge needle, followed by a 30-gauge needle. Three Lewis rats and three F344 rats at 8 weeks old were infected intraperitoneally (i.p.) with 2 × 10^6^ parasites of each strain and parasite viability of the inoculums was determined in a plaque assay after infection. At 60 days post-infection, the rats were sacrificed and the brains were harvested. Following homogenization of brains by passaging though a 21-gauge needle, cysts were stained by FITC-DBA. To detect the presence of parasite in the brains of infected rats, genomic DNA of homogenized brains was isolated using Qiagen DNeasy Blood & Tissue Kits (Qiagen). Diagnostic PCR targeting the B1 gene was performed by using the primer sets listed in **Supplementary Table 1**. To determine the anti-*Toxoplasma* IgG response of infected rats, serum was separated from the blood obtained at 60 days post-infection and anti*-Toxoplasma* IgG titer was detected using an enzyme-linked immunosorbent assay (ELISA). The plates were coated with 0.25 μg of whole parasite lysate produced by several freeze-thaw cycles. After blocking with 2% BSA in PBS-0.05% Tween-20, serial dilutions of serum was added and incubated at room temperature for at least 2 hours, followed by incubation with 1/2000 diluted HRP-conjugated goat anti-rat IgG at room temperature for 2 hours. Finally, after washing with PBS-0.05% Tween-20, 100 μl of substrate solution (ABTS solution from Sigma) was added to the wells and after 30 minutes the reaction was stopped by the addition of 50 μl of 0.3 M Oxalic acid, and optical density at 405 nm was measured. The titer corresponds to the dilution which gave an OD405 reading two-fold higher than the average of uninfected rat serum.

## Acknowledgements

This study was supported by the National Institutes of Health (R01-AI080621) awarded to J.P.J.S. K.M.C was supported by National Institutes of Health (F31-AI104170). M.A.H was supported by a Welcome Trust-MIT postdoctoral fellowship.

## Author contributions

J.P.J.S., Y.W. and K.M.C. designed experiments. Y.W. and K.M.C. performed and interpreted most of the experimental works. Y.W. performed all the experiments of figure 1, 2, 5, 7, 8, supplementary figure 1, 3 and 7. K.M.C. performed all the experiments of figure 3, 6, supplementary figure 4 and 8. Y.W. and K.M.C. performed all the experiments of figure 4, supplementary figure 2 and 5 Y.W. and L.O.S. conducted the *in vivo* infection experiment of figure 9. P.D.C.B. generated knockout strains and performed cell viability assay with these parasites in supplementary figure 6. M.A.H. and V.B. performed whole genomic sequencing and analyzed the sequencing data. P.P. and A.M. performed necropsy and pathological observation for *in vivo* studies. J.P.J.S., Y.W. and K.M.C. wrote the paper with contributions from all authors.

---

**Supplementary Table 1. Sequences of primers used in this study.**

HA-tag sequence is bolded. Restriction enzyme sites are underlined.

**Supplementary Figure 1. Neither Caspase-1 inhibitor nor NLRP3 inflammasome inhibitor affect parasite invasion**. (**A**) Lewis rat BMDMs (1×10^5^) were stimulated with or without 50 μM of VX765 or 10 μM of MCC950 for 2 hours followed by infection with 1×10^5^ of *Toxoplasma* type I (RH) parasites for 30 minutes. Quantification of the invading and invaded parasites per host nucleus. Data are displayed as the average values with scatterplots for each independent experiment (*n* = 3; error bars, ±SD; ns, not significant; one-way ANOVA with Kruskal-Wallis test). (**B**) Confluent HFFs were stimulated with or without VX765 (50 μM) or MCC950 (10 μM) followed by infection with parasites for 4 days. The area of at least 40 plaques per experiment was measured. Data are displayed as the average values with scatterplots for each independent experiment (*n* = 2; error bars, ±SD; ns, not significant; one-way ANOVA with Kruskal-Wallis test). (**C**) Lewis rat BMDMs primed with 100 ng/ml of LPS for 2 hours were treated with or without 10 μM of MCC950 for 2 hours followed by stimulating with or without 10μM of Nigericin for 2 hours. Macrophage viability was measured via MTS assay. Data are displayed as the paired scatterplots (*n* = 3; \**p* < 0.05, \*\**p* < 0.01, ns, not significant; student’s t-test). (**D**) IL-1β secretion was measured using ELISA on the cell supernatants from (C). Data are displayed as the paired scatterplots (*n* = 3; \**p* < 0.05, ns, not significant; student’s t-test).

**Supplementary Figure 2. PCR confirming knockout of candidate genes.** (**A**) Schematic diagram depicting the genomic loci of the genes of interest (GOI) (top) and the CRISPR/Cas9-targeting site (red box), linearized pTKOatt plasmid containing HXGPRT selection cassettes (middle) was used as repair template to disrupt GOI loci (bottom) after mycophenolic acid and xanthine selection. P1 and P2 refer to primers used for checking locus disruption. (**B**) Schematic diagram depicting the strategy used for making double/triple knockout. The *GRA42* or/and *GRA43* locus in Δ*gra35* parasites was disrupted by CRISPR/Cas9 cleavage and linearized pLoxp-DHFR-mCherry plasmid containing DHFR-TS selection cassette was used for NHEJ repair of the double stranded break. After pyrimethamine selection and limiting dilution, single clones with DHFR-mCherry integrated into the *GRA42* locus and an intact *GRA43* locus were used as the Δ*gra35*Δ*gra42* strain; single clones with an intact *GRA42* locus and DHFR-mCherry integrated into the *GRA43* locus were used as the Δ*gra35*Δ*gra43* strain; single clones with DHFR-mCherry integrated into both loci were used as triple knockout strain. (**C**) Genomic DNA was isolated from clones and used as template. Knockout was determined by failure to amplify the gene of interest using P1 and P2 as primers. DNA quality was assessed by amplifying *TGGT1_309160* (i and vi, for Δ*myr1* and single, double and triple knockout of *GRA35, GRA42* and *GRA43*), *B1* gene (ii, for Δ*TGGT1_248260*) or GRA35 (iii and iv, for Δ*TGGT1_203040* and Δ*rop17*).

**Supplementary Figure 3. Predicted structure and synonymous non-synonymous analysis of *GRA35, TGGT1_236870* and *TGGT1_237015***. (**A**) PSIPRED was used for secondary structure prediction (Jones 1999). Red stars indicate the mutation site in amino acid sequence of mutant #1, #2 and #3. Blue stars indicate the *GRA35* mutation site in amino acid sequence of mutant #4. Signal peptide, grey box; α-helices, green box. TmHMM2.0 was used for transmembrane domain (TM) prediction (Krogh et al. 2001). TM, blue box. Coiled-coil domain was analyzed using http://www.ch.embnet.org/software/COILS_form.html (Lupas, A., 1991). Coiled-coil region, yellow box. (**B, C and D**) SNAP was used for synonymous non-synonymous analysis (https://www.hiv.lanl.gov/content/sequence/SNAP/SNAP.html) (Korber B. 2000). Coding sequence of *GRA35* (**B**), *TGGT1236870* (**C**) and *TGGT1237015* (**D**) from 64 strains (ToxoDB 29 release) were analyzed using SNAP V2.1.1.

**Supplementary Figure 4. Mutant #4 does not induce cell death in Lewis rat macrophages.** Lewis rat BMDMs were infected with WT parasites or mutant clone #4, which was isolated from the pool of mutagenized parasites (MOI = 1) for 24 hours. Macrophage viability was measured via MTS assay. Data are displayed as the paired scatterplots (left, *n* = 4; \*\**p* < 0.01; student’s t-test). The right scatterplots are showing the cell viability difference between mutant #4 with WT parasites in each paired experiment. Horizontal bars represent the median cell viability difference.

**Supplementary Figure 5. GRA35, TGGT1_236870 and TGGT1_237015 have orthologues in *Hammondia, Neospora* and *Besnoitia***. Alignments of primary peptide sequences using PRALINE. (**A**) Alignment of GRA35 from Type I, II and III *Toxoplasma gondii, Hammondia hammondi* HHA_226380, *Neospora caninum* NCLIV_046580 and NCLIV_047520, *Besnoitia besnoiti* BESB_060230 and BESB_061290, and *Toxoplasma gondii* TGGT1_225160, GRA36 and TGGT1_257970. (**B**) Alignment of *Toxoplasma* TGGT1_236870, TGME49_236870 and TGVEG_236870, *Hammondia* HHA_236870, *Neospora* NCLIV_050780 and *Besnoitia besnoiti* BESB_036500. (**C**) Alignment of *Toxoplasma* TGGT1_237015, TGME49_237015 and TGVEG_237015, *Hammondia* HHA_237015 and *Neospora caninum* NCLIV_050915 and *Besnoitia besnoiti* BESB_036360.

**Supplementary Figure 6. *GRA35* gene family members are not involved in *Toxoplasma* induced cell death in Lewis rat BMDMs**. (**A**) Schematic diagram depicting the genomic loci of GOI (top) and the CRISPR/Cas9-targeting site, linearized pLoxp-DHFR-mCherry plasmid containing DHFR-TS selection cassettes (middle) was used as a repair template to disrupt GOI loci (bottom) after pyrimethamine selection. P1 and P2 refer to primers used for checking loci disruption; P1 and P3 refer to primers used for checking repair template integration. (**B**) PCR confirming individual knockout of *GRA35* gene family members with indicated primers. Genomic DNA isolated from each knockout parasites was used as template. (**C**) Lewis rat BMDMs were infected with WT parasites or the parasites in which *GRA35, TGGT1_225160, GRA36* or *TGGT1_257970* was knocked out (Δ*gra35*, Δ*TGGT1_225160*, Δ*gra36* or Δ*TGGT1_257970*) (MOI = 1) for 24 hours. Macrophage viability was measured via MTS assay. Data are displayed as the paired scatterplots (left, *n* = 3; all knockout strains vs. WT, \*\**p* < 0.01, ns, not significant; student’s t-test). The right scatterplots are showing the cell viability difference between indicated strains and WT parasites in each paired experiment. Horizontal bars represent the median cell viability difference.

**Supplementary Figure 7. GRA35, GRA42 or GRA43 mutants exhibit normal growth *in vitro*.** (**A**) Confluent Brown Norway rat immortalized fibroblasts infected with type II WT parasites (ME49-RFP) or the type II parasites in which *GRA35, GRA42* or *GRA43* was knocked out (ME49-RFPΔ*gra35*, ME49-RFPΔ*gra42* or ME49-RFPΔ*gra43*) for 7 days. The area of at least 40 plaques per experiment was measured. Data are displayed as the average values with scatterplots for each independent experiment (*n* = 3; error bars, ±SD; ns, not significant; one-way ANOVA with Kruskal-Wallis test). (**B**) F344 rat BMDMs were infected with WT parasites, parasites in which *GRA35, GRA42* or *GRA43* was knocked out (Δ*gra35*, Δ*gra42* or Δ*gra43*) or knockout parasites complemented with WT alleles of *GRA35, GRA42* or *GRA43* (Δ*gra35 + GRA35*, Δ*gra42 + GRA42* or Δ*gra43* + *GRA43*) (MOI = 1) for 24 hours. Macrophage viability was measured via MTS assay. Data are displayed as the grouped column combined with Lewis rat cell viability data showed at Figure 4C. Red indicates the average cell viability (+ SD) of Lewis rat BMDMs infected with indicated parasites; blue indicates the average cell viability (+ SD) of F344 rat BMDMs infected with indicated parasites (*n* = 3). (**C**) IFAs were performed on type II WT parasites (ME49-RFP) or the type II parasites in which *GRA35, GRA42* or *GRA43* was knocked out (ME49-RFPΔ*gra35*, ME49-RFPΔ*gra42* or ME49-RFPΔ*gra43*) as developing bradyzoite stages (for 3 days in alkaline pH 8.2). FITC-conjugated DBA was used to visualize the cyst wall. Images were taken at identical exposure times for each channel (scale bar = 10 μm). Image is representative of two independent experiments.

**Supplementary Figure 8. *Neospora caninum* is able to induce cell death in Lewis rat macrophages. Lewis BMDMs primed with LPS (100 ng/ml, 2 hours) or left untreated and infected with indicated parasites (*Toxoplasma*, RH; *Neospora*, NC-1) at MOI =1 for 24 hours.** (**A**) Cell viability was measured using an MTS assay. (**B**) IL-1β secretion was measured using ELISA on cell supernatants. Data shown are the average of two experiments, Error bars, +SD.

